# Spatio-temporal modelling of *Leishmania infantum* infection among domestic dogs: a simulation study and sensitivity analysis applied to rural Brazil

**DOI:** 10.1101/418558

**Authors:** Elizabeth Buckingham-Jeffery, Edward M. Hill, Samik Datta, Erin Dilger, Orin Courtenay

## Abstract

**Background:** The parasite *Leishmania infantum* causes zoonotic visceral leishmaniasis (VL), a potentially fatal vector-borne disease of canids and humans. Zoonotic VL poses a significant risk to public health, with regions of Latin America being particularly afflicted by the disease.

*Leishmania infantum* parasites are transmitted between hosts during blood feeding by infected female phlebotomine sand flies. With a principal reservoir host of *L. infantum* being domestic dogs, limiting prevalence in this reservoir may result in a reduced risk of infection for the human population. To this end, a primary focus of research efforts has been to understand disease transmission dynamics among dogs. One way this can be achieved is through the use of mathematical models.

**Methods:** We have developed a stochastic, spatial, individual-based mechanistic model of *L. infantum* transmission in domestic dogs. The model framework was applied to a rural Brazilian village setting with parameter values informed by fieldwork and laboratory data. To ensure household and sand fly populations were realistic, we statistically fit distributions for these entities to existing survey data. To identify the model parameters of highest importance, we performed a stochastic parameter sensitivity analysis of the prevalence of infection among dogs to the model parameters.

**Results:** We computed parametric distributions for the number of humans and animals per household and a non-parametric temporal profile for sand fly abundance. The stochastic parameter sensitivity analysis determined prevalence of *L. infantum* infection in dogs to be most strongly affected by the sand fly associated parameters and the proportion of immigrant dogs already infected with *L. infantum* parasites.

**Conclusions:** Establishing the model parameters with the highest sensitivity of average *L. infantum* infection prevalence in dogs to their variation helps motivate future data collection efforts focusing on these elements. Moreover, the proposed mechanistic modelling framework provides a foundation that can be expanded to explore spatial patterns of zoonotic VL in humans and to assess spatially targeted interventions.

## 1 Background

Zoonotic visceral leishmaniasis (VL) is a potentially fatal disease of humans and canids caused by the parasite *Leishmania infantum*. These parasites are transmitted between hosts during blood-feeding by infected female phlebotomine sand fly vectors [1, 2]. Zoonotic VL poses a significant risk to public health, being endemic in 65 countries in regions of Latin America, the Mediterranean, central and eastern Asia, and East Africa, with a case fatality rate of 90% in humans if left untreated [3–6].

Human infection has not been proven to be able to maintain *L. infantum* transmission without an infection reservoir [5]; the only proven reservoir host is domestic dogs [3–5]. Sand flies readily feed upon many other animal species, which act as important blood meal sources that support egg production. However, aside from domestic dogs these other animal species are considered “dead-end” hosts for parasite transmission since generally they do not support *Leishmania* infections and/or are not infectious. For most sand fly vector species, host preference is usually related to host biomass rather than to specific identity [7]. As a consequence, in addition to dogs and humans, domestic livestock living in close proximity to humans, such as chickens, pigs and cattle, are epidemiologically significant blood meal sources for sand flies [8, 9].

A primary focus of research efforts has been to understand the dynamics of *L. infantum* transmission among dogs, with the intent that limiting prevalence in this reservoir will result in a reduced risk of zoonotic VL infection for the human population. One way this can be achieved is through the use of mathematical models.

Mathematical models are a tool that allow us to project how infectious diseases may progress, show the likely outcome of outbreaks, and help to inform public health interventions. Through sand fly abundance and seasonality, *L. infantum* infection, and thus VL cases, has both spatial and temporal dependencies. There is, however, a surprising scarcity of mathematical models capable of capturing these spatio-temporal characteristics. A review by Rock et al. [10] found 24 papers addressing relevant modelling of VL, of which only two consider spatial aspects of transmission [11, 12]. Subsequent additions to the VL modelling literature since this review continue the tendency to exclude spatial heterogeneity in transmission. In particular, three recent studies (all published since the Rock et al. [10] review) have developed mathematical models that describe zoonotic VL dynamics in Brazil, but none contain any spatial aspects [13–15]. To our knowledge, there is presently no recorded work that specifies a spatial model of VL incorporating humans, vectors, reservoir hosts (dogs) and dead-end hosts.

One country severely afflicted by zoonotic VL is Brazil [6]. VL is endemic in particular regions of Brazil, exemplifying the spatial heterogeneity of the disease. In terms of canine VL, serological studies undertaken in endemic areas of Brazil have found prevalence of *L. infantum* infection to range from 25% [16] to more than 70% [17–20] depending on the diagnostic sample and test employed. A consequence of the burden of *L. infantum* infection in the canine reservoir is that Brazil has seen a steady rise in the number of human VL cases throughout the last 30 years [5, 21]. A reported 3,500 human VL cases occur in the country per year, 90% of all VL cases reported in the Americas [1, 3], with the actual incidence (allowing for under-reporting) estimated annually to be between 4,200 and 6,300 [1]. Accordingly, in Brazil importance is attached to the management of infection prevalence among domestic dogs to diminish the public health VL risk [22, 23].

To this end, we herein develop a novel spatio-temporal mechanistic modelling framework for *L. infantum* infection in domestic dogs. Applying the model to a rural Brazilian setting, we perform a sensitivity analysis to identify those model parameters that cause significant uncertainty in the predicted prevalence of *L. infantum* infection.

## 2 Methods

### 2.1 Model description

Informed by presently available field and laboratory data, we have developed a stochastic, spatial, individual-based, mechanistic model for *L. infantum* infection progression in domestic dogs in order to estimate *L. infantum* prevalence amongst the domestic dog population.

In brief, the model incorporates spatial variation of both hosts (adults and adolescents, children, dogs, and chickens) and vectors (sand flies) at the household level. Chickens represent dead-end hosts available to the sand fly vector; we do not refer explicitly to other dead-end hosts, such as pigs and cattle, as in the present study location chickens are the predominant domestic blood meal source for sand flies and chicken sheds yield the vast majority of sand flies captured within domestic areas [24–26]. Using a vectorial capacity type calculation, we derived a force of infection that gives the probability a dog will become infected with the *L. infantum* parasite via the sand fly vector. Infectious dogs increase the force of infection within a radius of their household. We tracked and reported as the output of the model the number of infected dogs each day.

Further details on each aspect of the model follow.

#### Households and hosts in space

We considered a configuration of rural households based on the latitude and longitude coordinates of 235 households in Calderao, a village on the island of Marajó in Northern Brazil (Figure 1). The household locations in Calderao are considered representative of a rural household spatial distribution in this endemic region. These household location data were collected as part of an epidemiological study of VL on Marajó between 2004 and 2005 where 99% of households were concurrently mapped by global positioning system technology (O. Courtenay and R.J. Quinnell, unpublished observations).

**Figure 1:**
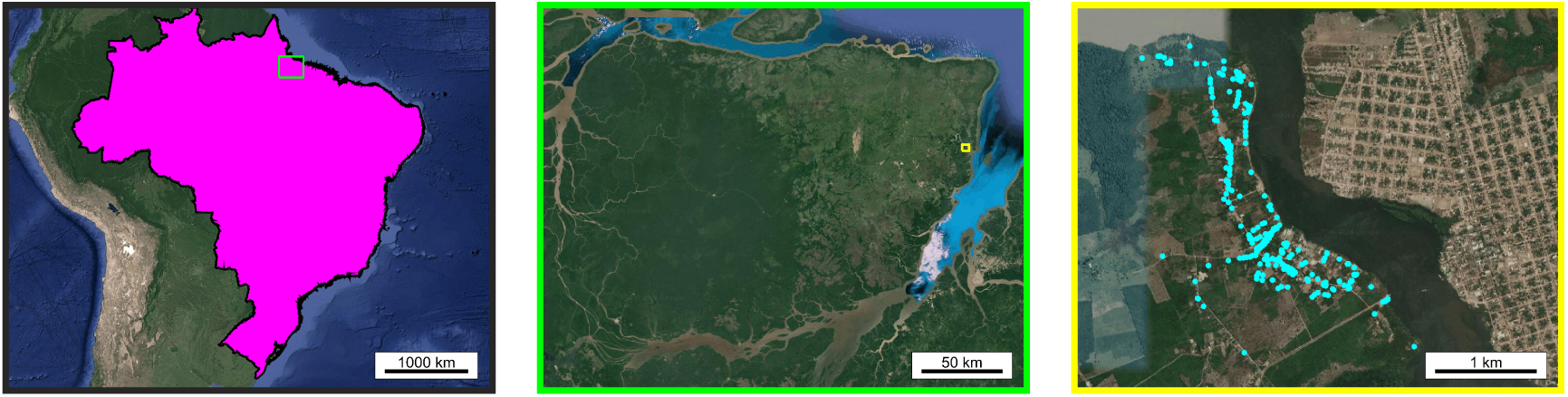
Locator maps. **(Left)** Map depicting Marajó, situated inside the light green box, within Brazil (shaded in magenta). **(Centre)** Map depicting Calderao village, situated inside the yellow box, within Marajó. **(Right)** Household locations within Calderao village (cyan filled circles). All map data are from Google and plotted in MATLAB®.

The number of each type of host at each household was assigned in each model run by sampling from distributions of host numbers per household (Figure 2). We obtained these distributions by fitting to survey data from the Marajó region collected in July and August of 2010 at 140 households across seven villages [27]. Further details of this data and obtaining the distributions can be found in Additional File 1.

**Figure 2:**
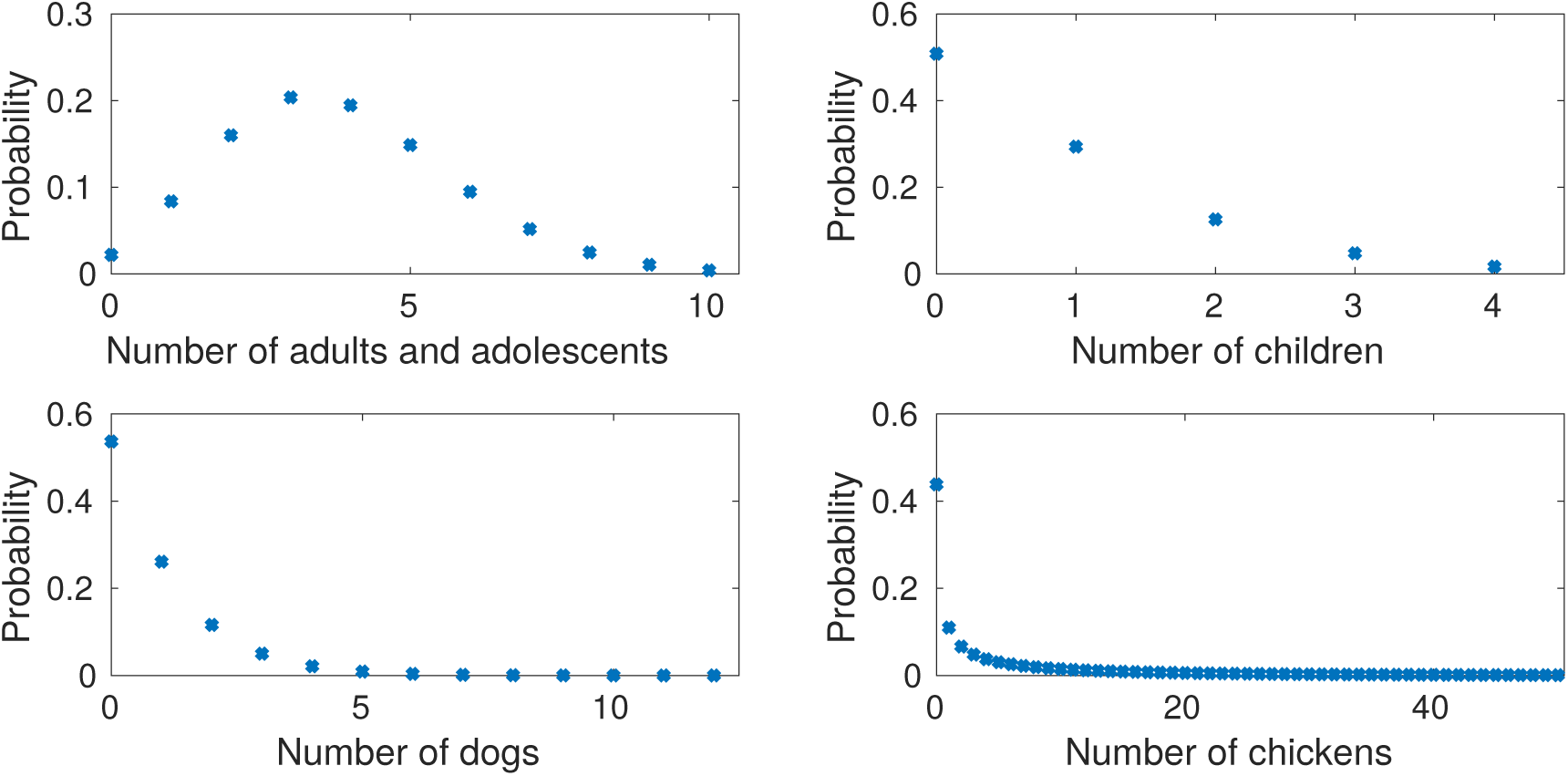
Distributions of the number of hosts per household. **(Top left)** Number of adults and adolescents; **(Top right)** children; **(Bottom left)** Number of dogs; **(Bottom right) Number of chickens.** Full details on how these distributions were obtained can be found in Additional File 1.

#### Infection progression in dogs

The natural history of *L. infantum* infection in dogs consists of susceptible and infected states. Prior work has established heterogeneities in the infectiousness of dogs (transmission of *L. infantum* to the vector) [2, 28, 29]. Specifically, this heterogeneity in infectiousness results in infected states representing highly infectious dogs (responsible for 80% or greater of all observed transmission events), mildly infectious dogs (contributing to 20% or less of total transmission events), and noninfectious dogs that, although infected, never transmit the *L. infantum* parasite back to susceptible sand flies [28].

For modelling purposes we therefore stratified infected dogs into four states: (i) latently infected; (ii) never infectious; (iii) low infectiousness; (iv) high infectiousness (Figure 3). Susceptible dogs became latently infected at a rate dependent on the force of infection; full details of this will follow. Movement between the latently infected state and the remaining three infected states occurred at constant rates. Note that a fully recovered state was not included as the complete cure of *L. infantum* infected dogs is rare (even after treatment), validated by experimental observations finding minimal seroreversion from *L. infantum* parasite seropositivity [30].

**Figure 3:**
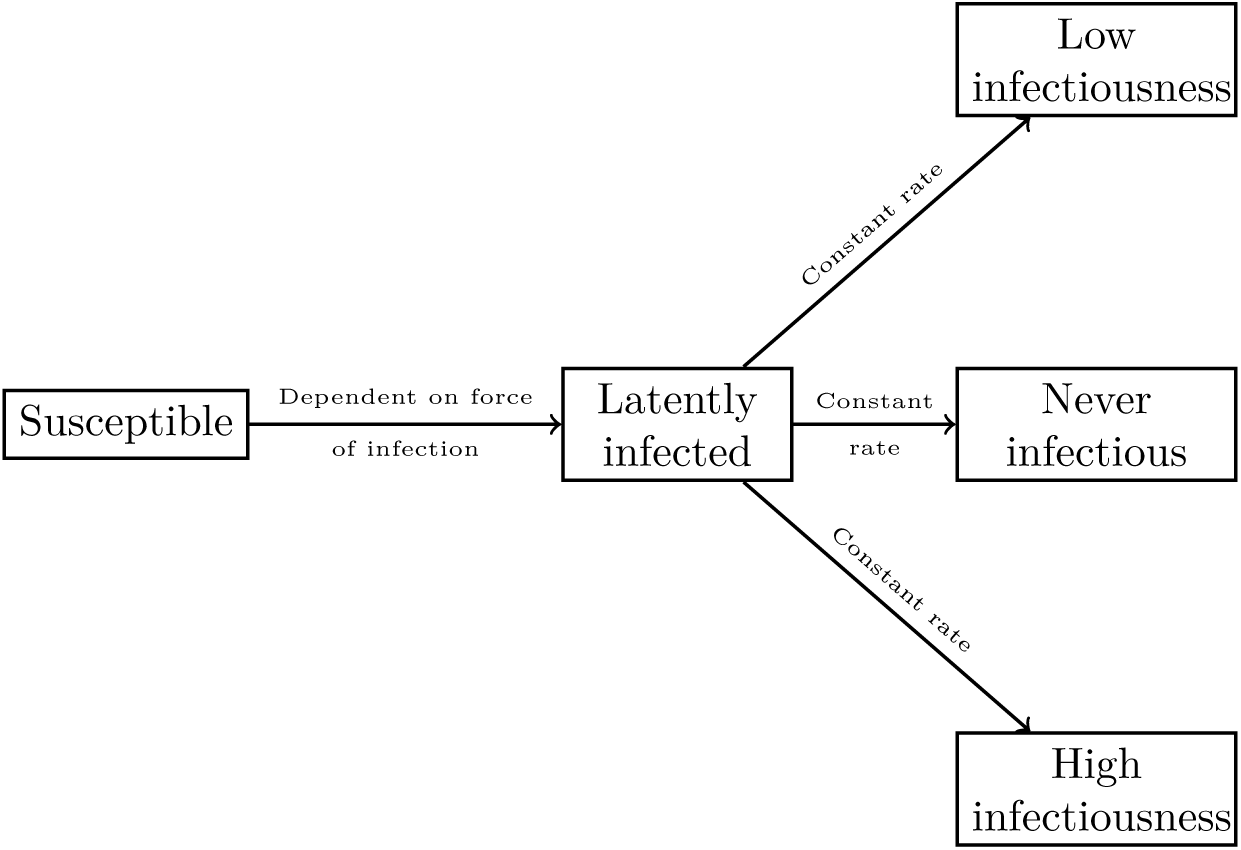
Model of *L. infantum* infection status in dogs. Death and replacement of deceased dogs (through birth and immigration) are not shown in the figure

Deaths could occur from every state in the model and the mortality rates differed between states. Upon death from any state, a new dog was introduced into the same household at a given replacement rate. Newly-introduced dogs were placed either in the susceptible state or one of the infected states, encapsulating both birth and immigration into the study region. It follows that the initial dog populations corresponded to the maximum attainable population size per household.

#### Force of infection

Sand fly dynamics operate on a faster time-scale compared to the other host species and processes considered in the model; sand flies have an estimated life expectancy of a number of weeks at most [10]. For that reason, we did not explicitly track the transitions of sand flies between the susceptible and infectious states at an individual level. We instead considered sand fly populations at each house as a collective which exert a force of infection, *λ*, on dogs at household *h* at time *t* in the following way,

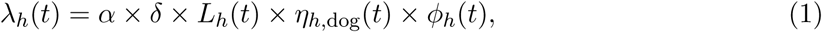

where *α* is the biting rate of sand flies, *δ* is the probability of *L. infantum* transmission to dogs as a result of a single bite from an infectious sand fly, *L*_*h*_ is the abundance of sand flies at household *h, η*_*h,*dog_ is the probability of sand flies biting dogs at household *h* as opposed to any other host, and *ϕ*_*h*_ is the proportion of sand flies that are infectious at household *h*.

As most sand fly activity occurs in the evening when the majority of hosts will be within their household [31, 32], we discretised our simulations into daily time steps. Using daily time steps gave the following probability for a susceptible dog at household *h* to become infected on day *t*:

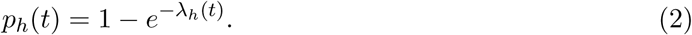

The biting rate and probability of an infected sand fly transmitting *L. infantum* to a dog as a result of a single bite were constant in the model. In contrast, sand fly abundance, host preference, and the proportion of sand flies infected at each household were time-dependent; we now outline the computation of each time-dependent component.

##### Sandfly abundance

Sand fly trapping data from villages in Marajó were used to obtain realistic estimates of the abundance of sand flies, *L*_*h*_, at households. As sand fly populations have been observed to exhibit temporal dependencies *L*_*h*_ comprised of two parts: a constant initial estimate and a seasonal scaling factor.

Data on the abundance of female sand flies, specifically the vector species *Lutzomyia longipalpis*, were available from a previous study of 180 households in fifteen villages on Marajó island where sand fly numbers were surveyed using CDC light-traps [24]. The constant initial estimate of abundance was sampled from these data and scaled by the expected proportion of unobserved female sand flies at households. Data on the mean number of female *Lutzomyia longipalpis* trapped over an eight month period across eight different households in the village of Boa Vista, Marajó [33] were then used to find the seasonal scaling factor. Full details of this procedure to estimate sand fly abundance can be found in Additional File 1.

##### Host preference

To parameterise sand fly biting preference towards the host species of interest, we drew on findings from field and laboratory experiments performed in this setting by Quinnell et al [7]. These experiments concluded that the attractiveness of the three host species we consider (humans, dogs and chickens) to the *Lutzomyia longipalpis* vector seemed to largely be a function of the relative host sizes.

These experimental findings were used to allocate a portion of sand fly bites to each host type at each household, via each host type being assigned the following biomass value relative to chickens:

- 1 dog = 2 chickens,
- 1 child = 5 chickens,
- 1 adult or adolescent = 10 chickens (using adult-child ratio: 1 adult = 2 children).

The preference, *η*_*h,x*_, towards host type *x* at household *h* was computed as a simple proportion of the total biomass,

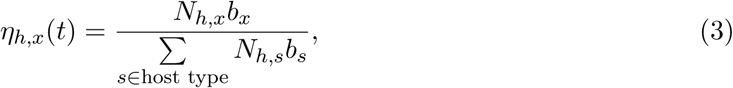

where *N*_*h,x*_ is the number of host type *x* at household *h* and *b*_*x*_ is the biomass of host type *x* relative to chickens. So, for example, *b*_dog_ = 2.

##### Proportion of infectious sand flies

The proportion of infectious vectors at household *h* was comprised of a time-independent background level of prevalence that was constant across all households and an additional proportion dependent on the number of infectious dogs in the neighbourhood of household *h*. We informed the radius defining this neighbourhood by matching it to the maximum sand fly travel distance (taken as 300m at the baseline with a range from 20m to 2km to fully explore the parameter space [34], see Table 1). The contribution from each type of infectious dog (high and low infectiousness) was computed separately under an assumption that 80% of transmission from dogs to sand flies is caused by high infectiousness dogs, with the remaining 20% of total transmission events contributed by infected dogs with low infectiousness [28]. Further details on our calculation of the proportion of sand flies that were infectious are given in Additional File 1.

**Table 1:**
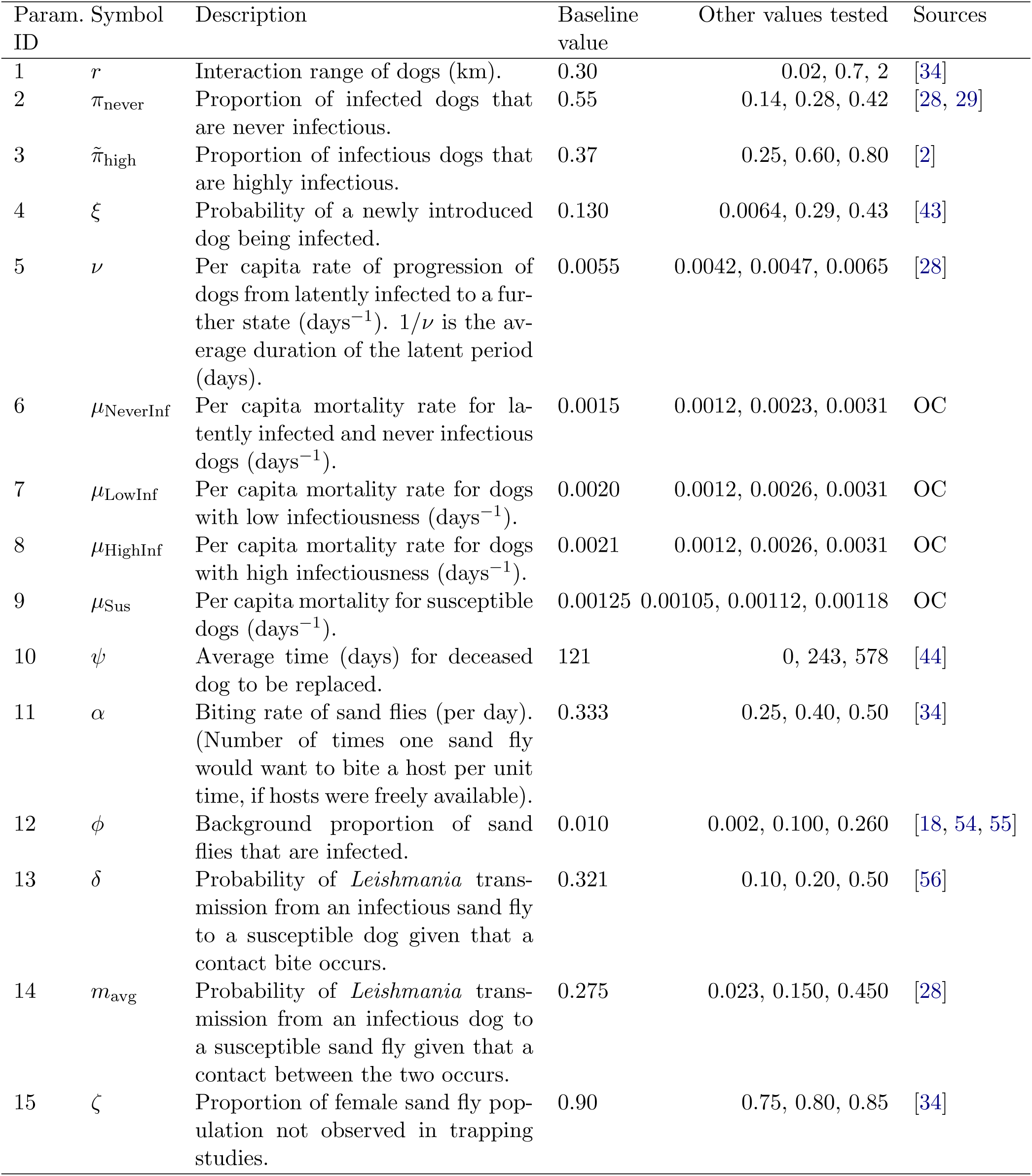
Description of measurable biological variables that are used to inform parameters (either directly or after performing additional calculations) in the model. Source listed as OC denotes (O. Courtenay, unpublished observations).

### 2.2 Model outputs

Being a stochastic model, the infection dynamics vary on separate simulation runs even with all parameters and other model inputs remaining fixed. By running the model multiple times we obtain an ensemble of model outputs. This collection of model outputs permits the calculation of a variety of summary statistics describing the epidemiology of *L. infantum* infection among domestic dogs, such as prevalence and incidence.

We focus here on the prevalence of infection. To clarify, an infection case refers to any dog harbouring *L. infantum* parasites, including those with and without canine VL symptoms. Thus, we defined infection prevalence at time *t* as the aggregated percentage of dogs in the latently infected, never infectious, low infectiousness and high infectiousness states, which is equivalent to calculating the proportion of dogs not in the susceptible state:

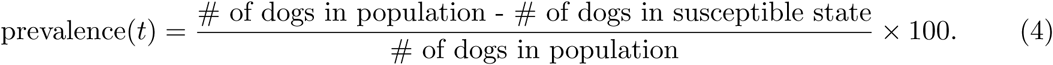

The daily prevalence estimates were used to obtain an average prevalence, defined as the mean of the daily prevalence estimates in a specified time period. Throughout this work, all average prevalence values were computed from the daily prevalence values over the final year (365 days) of each simulation run. Mathematically, with *T* denoting maximum time, average infection prevalence may be expressed as

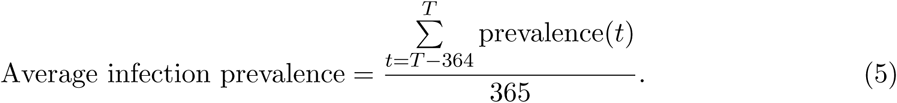

### 2.3 Model summary

In summary, the arrangement of and interaction between the individual pieces of our stochastic, spatial, individual-based model for *L. infantum* infection dynamics in dogs are displayed in Figure 4. We refer to the process in Figure 4 as one run of the simulation.

**Figure 4:**
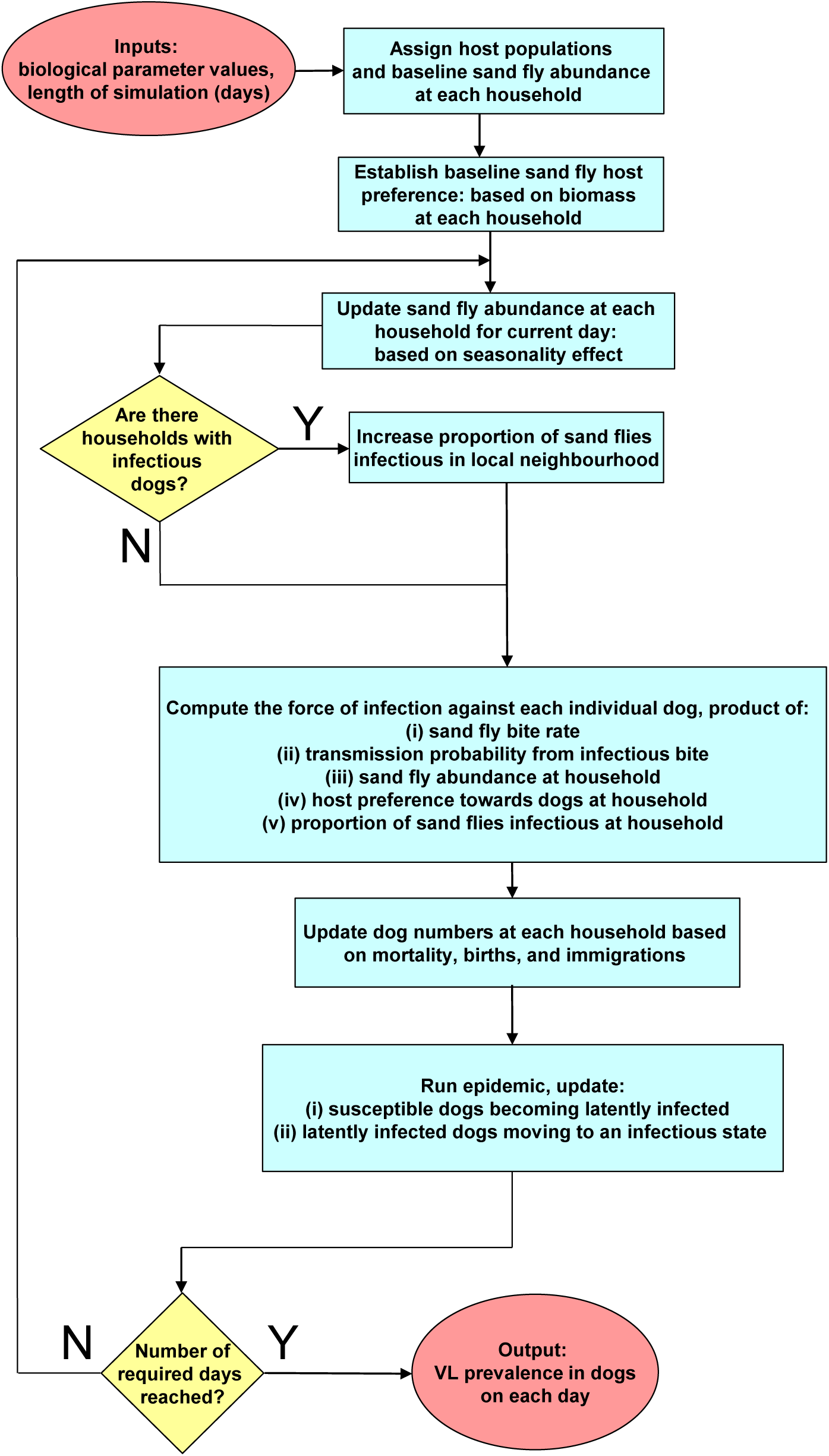
Visual schematic of model framework for each simulation run. Red filled ovals represent model inputs and outputs; blue filled rectangles represent actions; yellow filled diamonds represent decisions.

### 2.4 Sensitivity analysis

#### Parameter values

We carried out a sensitivity analysis to determine the robustness of the model behaviour to the biological parameter values and to ascertain which parameters had a high impact on the average prevalence as predicted by the model. The values tested for each parameter were within plausible ranges informed via published estimates from the literature and unpublished fieldwork data (Table 1).

We undertook a one-at-a-time sensitivity analysis. That is, each parameter was varied in turn while all others remained at their baseline value. We considered 46 parameter sets (Table 1), and for each individual parameter set we performed 1000 separate model simulation runs. The elapsed simulation time in each run corresponded to ten years.

#### Sensitivity coefficients

In addition to comparing the changes in average prevalence given by each parameter set, we computed sensitivity coefficients. These reflect the ratios between the size of the change in a model output (in this case, the change in average VL prevalence) with the corresponding size of the change in the parameter [35]. The sensitivity coefficients therefore account for the different ranges in the values tested for each parameter (Table 1) and ensure that the parameters can be sensibly compared.

However, in a stochastic modelling framework, such as this one, model outputs do not take a unique value. To account for stochastic fluctuations while still allowing us to critically analyse the sensitivity of the model parameters, we therefore calculated stochastic sensitivity coefficients (as outlined in Damiani et al. [36], comprehensive explanation in Additional File 1). We ranked the parameters according to the stochastic sensitivity coefficients, with a larger sensitivity coefficient corresponding to a parameter with higher sensitivity of average VL prevalence to its variation.

All simulations were performed in MATLAB® versions R2014a to R2015a. All other computations and plots were carried out in MATLAB® version R2016b or later.

## 3 Results

### 3.1 Model simulations - Baseline parameters

As a form of model validation, we checked the plausibility of infection prevalence predictions while each biological parameter was fixed to its baseline value (Table 1). Under these baseline parameter values, the daily prevalence in dogs was generally between 46% and 68%. Averaging over 1000 separate model simulation runs, the median trace for daily prevalence in dogs lay between 55% and 59%. Seasonal oscillations in the median prevalence remained observable across time, though ordinarily less pronounced compared to the seasonality-induced changes in prevalence apparent in a single simulation run (Figure 5).

**Figure 5:**
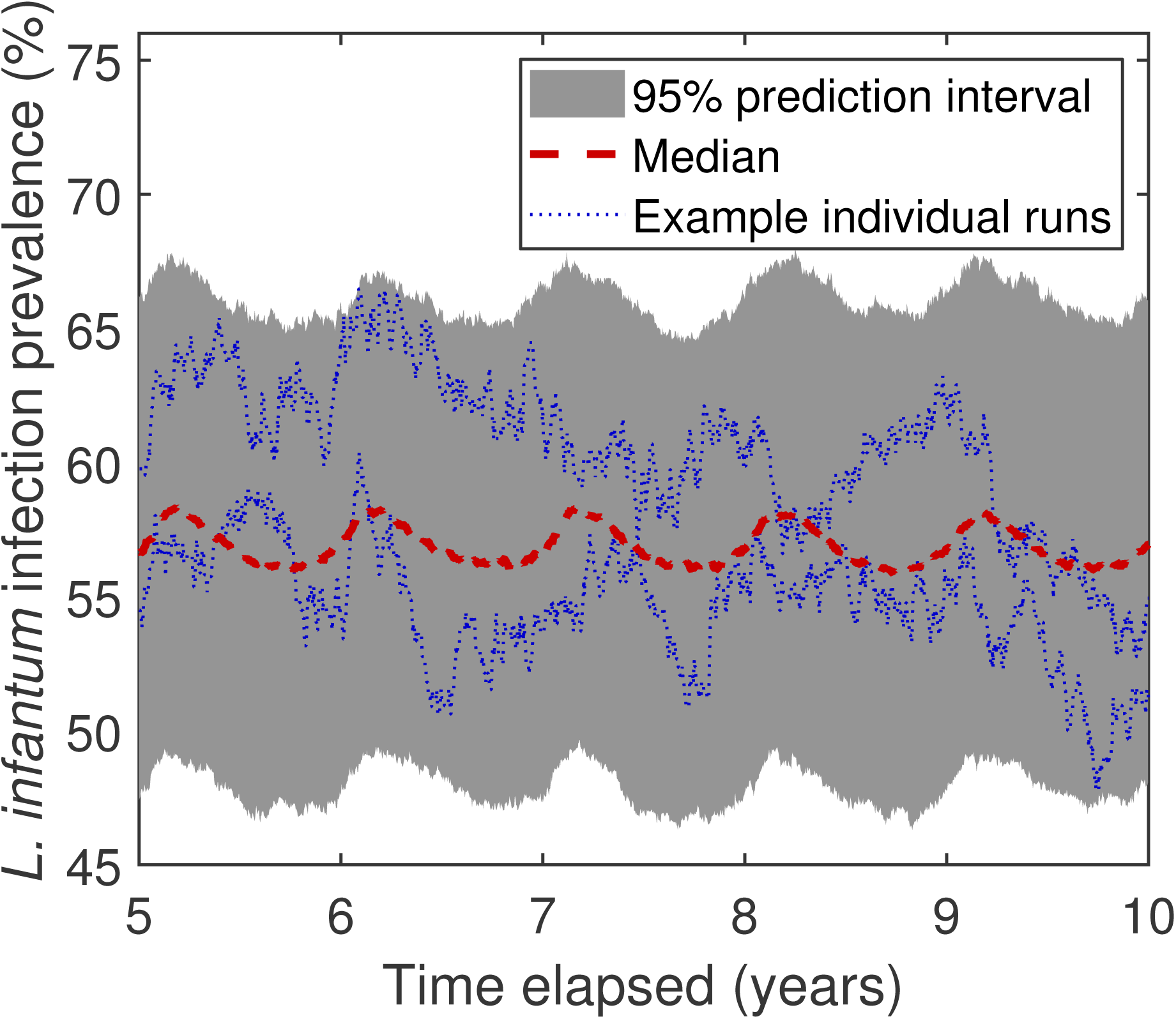
Simulated daily prevalence in domestic dogs using baseline biological parameters. Dashed, red line corresponds to the median prevalence and the grey, filled region depicts the 95% prediction interval at each timestep obtained from 1000 simulation runs. Blue, dotted lines correspond to measured prevalence from two individual simulation runs.

#### 3.2. Sensitivity analysis

Under baseline parameter values, the median of the average infection prevalence over 1000 simulation runs was 57% (95% prediction interval: [49%, 66%]). In addition, the ranges of the average infection prevalence distributions were quantitatively similar irrespective of the parameter set tested (Figure 6).

**Figure 6:**
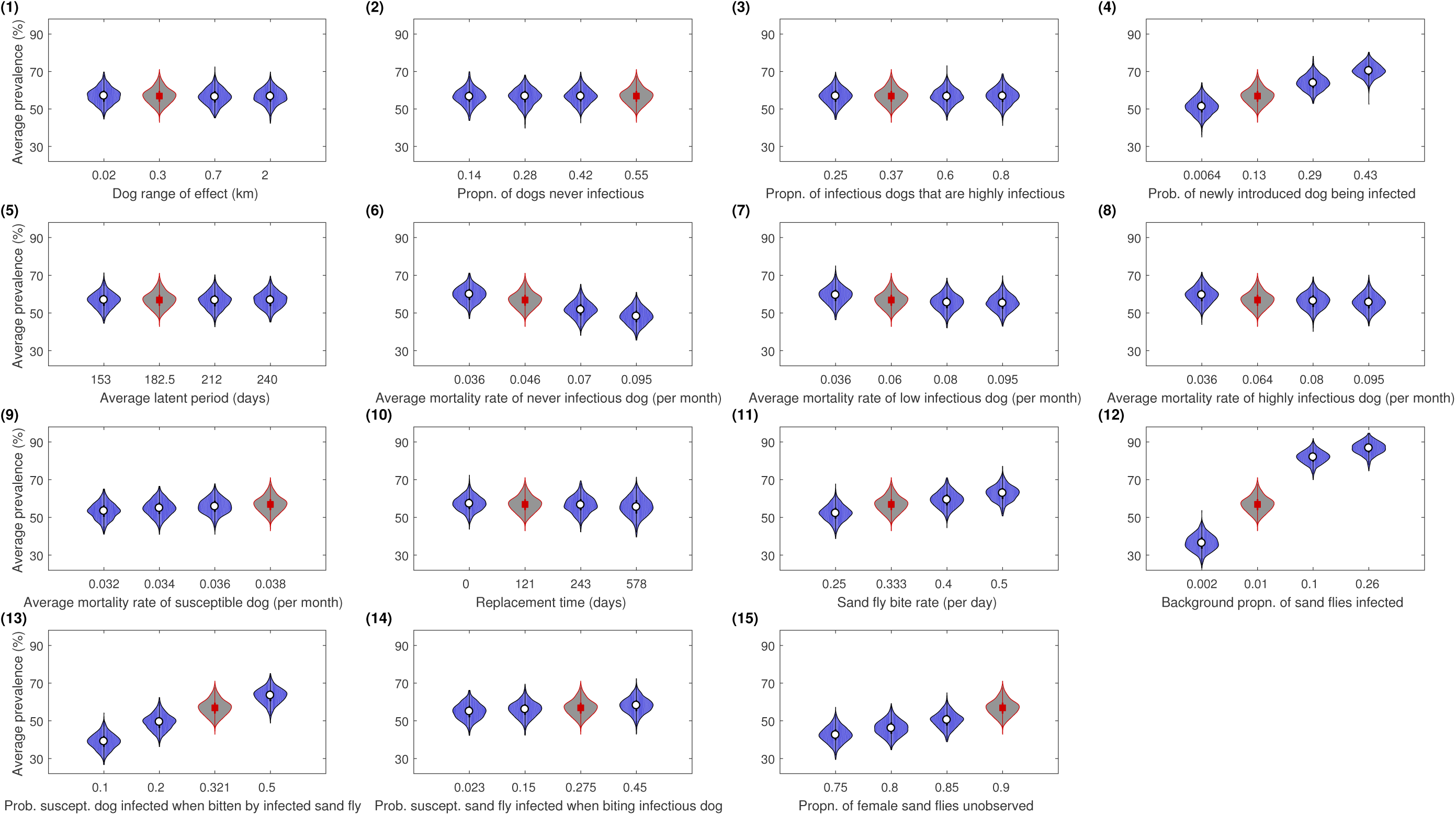
Violin plots for average infection prevalence under each biological parameter set. Panel numbering aligns with the parameter ID numbers in Table 1. The average infection prevalence was calculated by the daily prevalence values over the final year of each simulation run. For each parameter set, predicted average infection prevalence distributions were acquired from 1000 simulation runs. The violin plot outlines illustrate kernel probability density, i.e. the width of the shaded area represents the proportion of the data located there. For parameter sets corresponding to the use of the baseline parameter set, violin plot regions are shaded grey with estimated median values represented by a red square. In all other instances, violin plot regions are shaded blue with median values depicted by a white circle.

Among the 46 parameter sets tested, the largest median average infection prevalence prediction (87%) was obtained when the background proportion of sand flies infected (parameter ID 12) was increased from its baseline value of 0.01 to 0.26 (with all other biological parameters fixed at baseline values). Similarly, the smallest median average infection prevalence prediction (36%) was obtained when the background proportion of sand flies infected was lowered to 0.002 (with all other biological parameters again fixed at baseline values). As a consequence, this parameter set had an approximate 50% shift in absolute value of the median across the range of tested values: the highest among the 15 biological parameters in this sensitivity analysis (Figure 6, panel (12)).

Moreover, when comparing the respective sensitivity test values in three other sand fly-associated parameter sets, sand fly bite rate (parameter ID 11), probability of a susceptible dog becoming infected when bitten by an infected sand fly (parameter ID 13) and proportion of female sand flies unobserved (parameter ID 15), in each case we found the median average infection prevalence to differ by over 10% across the range of values tested (Figure 6, panels (11,13,15)).

In the biological parameters associated with dogs, a visible rise in average infection prevalence was evident for parameter ID 4, the probability of a newly introduced dog being infected (Figure 6, panel (4)). On the other hand, for the average mortality rate of a never infectious dog (parameter ID 6), we saw a decrease of over 10% in the median estimates for average infection prevalence across the four tested values.

In all remaining parameter sets, the differences between the four median estimates for average infection prevalence were below 10% (Figure 6).

#### Parameter sensitivity rankings

By computing stochastic sensitivity coefficients and ranking the parameters by this measure, we discerned that the average infection prevalence was most sensitive to the probability of a newly introduced dog being infected (parameter ID 4). Of the four parameters linked to dog mortality (parameter IDs 6-9), the most critical was the mortality rate of never infectious dogs (parameter ID 6), which out of all 15 biological parameters under consideration ranked fourth overall (Figure 7).

**Figure 7:**
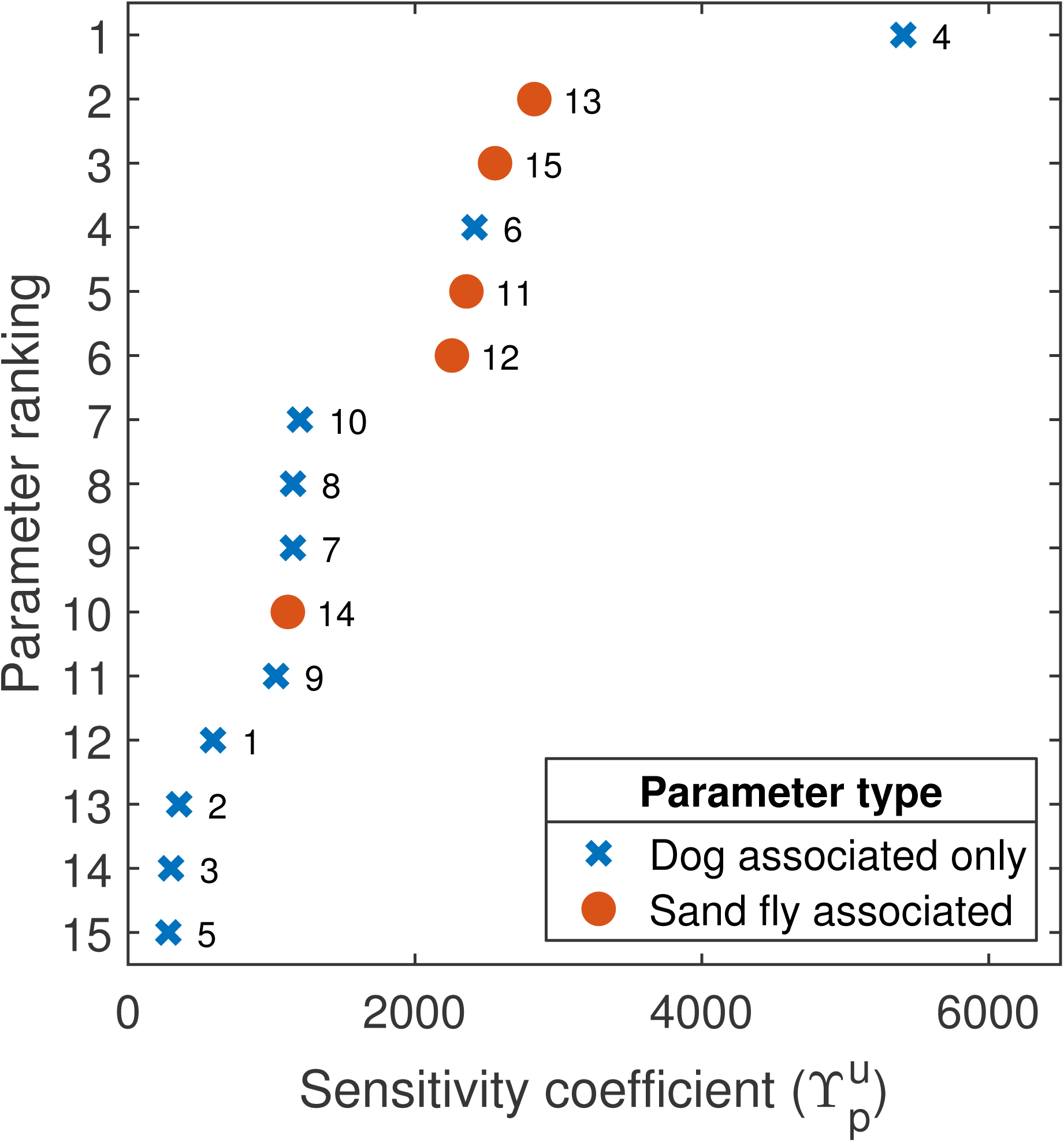
Stochastic sensitivity coefficient parameter ranking. The parameter ID linked to each stochastic sensitivity coefficient is placed aside the data point. Blue crosses denote those biological parameters associated with dogs. Filled orange circles correspond to biological parameters associated with sand flies. Average infection prevalence was most sensitive to parameter ID 4 (probability of a newly introduced dog being infected).

Four parameters associated with sand flies were among the top six most sensitive parameters (Figure 7). The only sand fly-associated parameter that was not among these top six most sensitive parameters was the probability of a susceptible sand fly becoming infected when biting an infectious dog (parameter ID 14).

## 4 Discussion

Despite zoonotic VL being spatially heterogeneous, there remains few spatially explicit mathematical models of *Leishmania* transmission to help inform infection and VL disease monitoring, surveillance and intervention efforts [10–12]. Amongst prior work, Hartemink et al. [11] predicted spatial sand fly abundance in southwest France to inform the construction of a basic reproductive ratio map for canine VL. However, these risk maps relied on sand fly abundance estimates from a single sampling timepoint; no temporal dynamics of sand fly abundance, and therefore of infection prevalence, were considered. A model developed by ELmojtaba et al. [12] was used to analyse whether a hypothetical human VL vaccination could successfully reduce prevalence when there is immigration of infected individuals into the population. While the model includes spatial aspects through the immigration mechanism, it lacks any explicit spatial structure in the modelled population.

In contrast, our study presents a novel spatio-temporal mechanistic modelling framework for *Leishmania* infection dynamics, incorporating humans, vectors, reservoir hosts (dogs) and dead-end hosts (chickens in this study; our nominal dead-end host species). We apply this model to a rural village setting based on empirical datasets measured on Marajó in Brazil to draw attention to those model inputs that cause significant uncertainty in the predicted prevalence of *L. infantum* parasites in domestic dogs.

### Curation of data

An integral part of the model set up involves incorporating data on host numbers per household, spatial sand fly abundances, and the temporal profile of sand fly abundances. The scarcity of exhaustive information on these population-level attributes necessitated that we fit distributions and smooth trend lines to small but informative datasets. The fitted host number distributions and sand fly abundance profiles offer a resource that may readily be applied in settings with similar social, environmental and climatic conditions.

### Sensitivity of *L. infantum* infection to biological parameter variation

Running model simulations using baseline biological parameter values set within plausible ranges determined from the literature generated infection prevalence predictions that were within the range of empirical estimates from endemic regions of Brazil [16–20]. Variation in infection estimates is expected as ultimately their precision depends on the type of diagnostic test used (e.g. molecular vs. immunological), diagnostic test sensitivity and specificity, the choice of clinical sample, and the stage of infection progression [17, 19, 20, 37]. Thus, for example, as dogs acquire parasitological infection prior to detection of serum containing anti-*Leishmania* specific antibodies (seroconversion), seroprevalence data may underestimate true infection rates.

The sensitivity parameter ranking reveals that ensuring sand fly vector associated parameters are well-informed warrants major attention; four out of the five parameters associated with sand flies were among the parameters with the highest sensitivity of average prevalence to their variation. Particularly sensitive were the parameters for the probability of transmission of infection from an infectious sand fly to a susceptible dog given that a contact between the two occurs (parameter ID 13) and the proportion of female sand flies not observed in trapping studies (parameter ID 15). It is unsurprising that the latter parameter displays high sensitivity; the proportion of female sand flies not observed in trapping studies directly affects the estimated sand fly abundance and thus the magnitude of the force of infection.

Ultimately, VL being a vector-borne disease means that infection events are driven by sand fly biting behaviour and sand fly interactions with hosts. Accordingly, finding greater sensitivity on infection prevalence when altering the parameters related to sand fly dynamics versus the majority of parameters conditioned solely on dogs is not unexpected and is in agreement with prior studies displaying the sensitivity of *Leishmania* transmission models to sand fly parameter values [13, 38]. Furthermore, the importance of understanding sand fly biology and biting behaviours in relation to transmission probability and control has been underpinned by laboratory experiments and observations in nature [32, 39–42].

Overall, the parameter with the highest sensitivity coefficient was the probability of a newly introduced dog being infected (parameter ID 4). Thus, reliably informing the relative amount of dog immigration into a region versus birth, plus the proportion of immigrant dogs already harbouring *L. infantum* parasites, is integral to providing reliable infection prevalence estimates. Studies of domestic dog migration are few, but in most dog populations losses and replacements appear relatively stable with estimates from Brazil of the percentage of new dogs being immigrants ranging from 37% to 50%, with up to 15% of immigrant dogs being *Leishmania* seropositive on arrival [43–45]. Given the heterogeneities in sand fly abundance and infection [42], even in highly endemic regions such as Marajó, migration of infected dogs between villages can have a significant impact on transmission as demonstrated here.

### Study limitations

Developing and parameterising an original mathematical framework in the face of limited data has its restrictions. First, we acknowledge that our findings are likely to be sensitive to the biomass-linked assumption for sand fly biting preference towards host species. The literature used to inform this assumption in the current model [7] is appropriate as it was conducted at the same site where most of the data used in the model were generated and is, we believe, the only experimental study of its type. However, the effect of alternative choices merits further investigation in tandem with field work for further data collection. Second, our analysis has focused on a single, rural household spatial configuration, although the selected configuration was chosen as representative of a typical village in Marajó, from where the majority of the parameter estimates were measured. Applying a similar methodological approach to semi-urban and urban populations would be informative and timely as zoonotic VL has recently expanded its geographical distribution to include urbanised communities [3, 46]. Such analysis offers the opportunity to quantify the impact of household spatial configuration on infection prevalence in domestic dogs across a range of environmental settings and the extent to which transmission is driven by the level of clustering or regularity in household locations. Finally, we assumed a maximum attainable dog population size per household and constant population sizes of other hosts. It would be of interest to explore the impact on infection prevalence among domestic dogs if there were to be an influx of alternative host livestock in close vicinity to households as dead-end host abundance is variably associated with infection risk [47–49].

### Further work

We anticipate this modelling framework being extended in a variety of ways. One future development would be to explore spatial patterns of zoonotic VL in humans resulting from the spatial distribution of *L. infantum* infection in domestic dogs. Our mechanistic approach for evaluating the force of infection is advantageous in that Equation (1) may be easily generalised to cater for host types other than dogs. Furthermore, while we considered a solitary dead-end host type, chickens, additional dead-end host types could seamlessly be incorporated using our modelling framework, allowing it to be used in settings where multiple livestock species are present.

Another application is to assist in intervention planning, where there is a need to employ the use of spatial models to predict best practice deployment of proposed controls through time and space. The spatial nature of our model makes it amenable to incorporating innovative, spatially-targeted vector and/or reservoir host control strategies that existing models were not designed to explore. One example, whose deployment nature is inherently spatial, is a pheromone-insecticide combination as a “lure and kill” vector control tool. Containing a long-lasting lure that releases a synthetic male sex pheromone, attractive to both sexes of the target sand fly vector [50, 51], this technology could be applied by disease control agencies to attract sand flies away from feeding on people and their animals and towards insecticide treated surfaces where they can be killed [50, 52]. To evaluate the impact of a pheromone lure via simulation, the intrinsic properties of the lure, such as its longevity and the radius within which it has an effect, necessitate the use of a spatio-temporal modelling framework such as the one presented here. A second example is the use of deltamethrin-impregnated dog collars which aim to protect dogs from sand fly bites [53]. Due to the decay of the effectiveness of the collars with time [53] and the spatial distribution of dogs in villages in Marajó, the effectiveness of this collar-based intervention could again be evaluated by our spatio-temporal modelling framework. With all repellent interventions, one must be careful to ensure that sand fly feeding is not diverted onto other hosts, including humans; an extended model variant considering zoonotic VL in humans could be used to estimate the size of this effect.

## 5 Conclusions

Zoonotic VL, caused by *Leishmania* parasites, is spatially heterogeneous and it is essential that monitoring, surveillance and intervention strategies take this variation into account. At the time of writing, there is a lack of spatially explicit mathematical models encapsulating *Leishmania* infection dynamics. We have developed a novel individual-based, spatio-temporal mechanistic modelling framework which, when parameterised according to data gathered from Marajó in Brazil, generated plausible *L. infantum* infection prevalence estimates.

Our study determined infection prevalence in dogs to be most strongly affected by sand fly associated parameters and the proportion of newly introduced (immigrant) dogs already infected. Identifying the biological factors with the greatest influence on expected infection prevalence motivates future data collection efforts into these particular elements; ensuring they are reliably informed will reduce the amount of uncertainty surrounding mathematical model generated predictions. Additionally, our mechanistic modelling framework provides a platform which can be built upon to further explore the spatial epidemiology of zoonotic VL in humans and to assess spatially-targeted interventions to inform VL response protocols.

## Supporting information

## Declarations

### Acknowledgements

The authors thank Deirdre Hollingsworth and Lloyd Chapman for helpful discussions, Gordon Hamilton for his role in funding acquisition, and Rupert Quinnell for provision of sand fly data. We acknowledge Chris Davies, Trystan Leng, Sophie Meakin, Tim Pollington and Emma Southall for their helpful feedback on the manuscript. The work utilised Queen Mary’s Mid-plus computational facilities supported by QMUL Research-IT and funded by Engineering and Physical Sciences Research Council grant EP/K000128/1.

## Author Contributions

**Conceptualisation:** OC

**Data Curation:** ED, OC

**Formal analysis:** EBJ, EH

**Funding Acquisition:** OC

**Investigation:** EBJ, EH

**Methodology:** EBJ, EH, SD

**Resources:** OC, ED

**Software:** EBJ, EH

**Supervision:** OC, ED

**Validation:** EBJ, EH

**Visualisation:** EBJ, EH

**Writing - original draft:** EBJ, EH

**Writing - review & editing:** EBJ, EH, SD, ED, OC

All authors read and approved the final version of the manuscript.

## Data availability

Parameter values used during this study are included in this article (Table 1). Code developed for the current study are available at https://github.com/EBucksJeff/VL_spatial_model. The raw datasets used and analysed during the current study are available from the authors on reasonable request and for use in the context of the research study.

## References

[1] Ready, P.: Epidemiology of visceral leishmaniasis. Clin. Epidemiol. 6, 147–154 (2014). doi:10.2147/CLEP.S44267

[2] Courtenay, O., Carson, C., Calvo-Bado, L., Garcez, L.M., Quinnell, R.J.: Heterogeneities in Leishmania infantum Infection: Using Skin Parasite Burdens to Identify Highly Infectious Dogs. PLoS Negl. Trop. Dis. 8(1), 2583 (2014). doi:10.1371/journal.pntd.0002583

[3] Martins-Melo, F.R., Lima, M.d.S., Ramos, A.N., Alencar, C.H., Heukelbach, J.: Mortality and Case Fatality Due to Visceral Leishmaniasis in Brazil: A Nationwide Analysis of Epidemiology, Trends and Spatial Patterns. PLoS One 9(4), 93770 (2014). doi:10.1371/journal.pone.0093770

[4] Vilas, V.J.D.R., Maia-Elkhoury, A.N.S., Yadon, Z.E., Cosivi, O., Sanchez-Vazquez, M.J.: Visceral leishmaniasis: a One Health approach. Vet. Rec. 175(2), 42–44 (2014). doi:10.1136/vr.g4378

[5] Conti, R.V., Moura Lane, V.F., Montebello, L., Pinto Junior, V.L.: Visceral leishmaniasis epidemiologic evolution in timeframes, based on demographic changes and scientific achievements in Brazil. J. Vector Borne Dis. 53(2), 99–104 (2016)

[6] World Health Organisation: Leishmaniasis - Fact Sheet (2018). http://www.who.int/en/news-room/fact-sheets/detail/leishmaniasis Accessed 27 Nov 2018.

[7] Quinnell, R.J., Dye, C., Shaw, J.J.: Host preferences of the phlebotomine sandfly Lutzomyia longipalpis in Amazonian Brazil. Med. Vet. Entomol. 6(3), 195–200 (1992)

[8] Morrison, A.C., Ferro, C., Tesh, R.B.: Host preferences of the sand fly lutzomyia longipalpis at an endemic focus of american visceral leishmaniasis in colombia. Am. J. Trop. Med. Hyg. 49(1), 68–75 (1993)

[9] Macedo-Silva, V.P. and Martins, D.R. and De Queiroz, P.V. and Pinheiro, M.P. and Freire, C.C. and Queiroz, J.W. and Dupnik, K.M. and Pearson, R.D. and Wilson, M.E. and Jeronimo, S.M. and Ximenes, M. de F.: Feeding preferences of Lutzomyia longipalpis(Diptera: Psychodidae), the sand fly vector, for Leishmania infantum (Kinetoplastida: Trypanosomatidae). J. Med. Entomol. 51(1), 237–244 (2014)

[10] Rock, K.S., le Rutte, E.A., de Vlas, S.J., Adams, E.R., Medley, G.F., Hollingsworth, T.D.: Uniting mathematics and biology for control of visceral leishmaniasis. Trends Parasitol. 31(6), 251–259 (2015). doi:10.1016/j.pt.2015.03.007

[11] Hartemink, N., Vanwambeke, S.O., Heesterbeek, H., Rogers, D., Morley, D., Pesson, B., Davies, C., Mahamdallie, S., Ready, P.: Integrated Mapping of Establishment Risk for Emerging Vector-Borne Infections: A Case Study of Canine Leishmaniasis in Southwest France. PLoS One 6(8), 20817 (2011). doi:10.1371/journal.pone.0020817

[12] ELmojtaba, I.M., Mugisha, J.Y.T., Hashim, M.H.A.: Vaccination model for visceral leishmaniasis with infective immigrants. Math. Methods Appl. Sci. 36(2), 216–226 (2013). doi:10.1002/mma.2589

[13] Sevá, A.P., Ovallos, F.G., Amaku, M., Carrillo, E., Moreno, J., Galati, E.A.B., Lopes, E.G., Soares, R.M., Ferreira, F.: Canine-Based Strategies for Prevention and Control of Visceral Leishmaniasis in Brazil. PLoS One 11(7), 0160058 (2016). doi:10.1371/journal.pone.0160058

[14] Zhao, S., Kuang, Y., Wu, C.-H., Ben-Arieh, D., Ramalho-Ortigao, M., Bi, K.: Zoonotic visceral leishmaniasis transmission: modeling, backward bifurcation, and optimal control. J. Math. Biol. 73(6-7), 1525–1560 (2016). doi:10.1007/s00285-016-0999-z

[15] Shimozako, H.J., Wu, J., Massad, E.: Mathematical modelling for Zoonotic Visceral Leishmaniasis dynamics: A new analysis considering updated parameters and notified human Brazilian data. Infect. Dis. Model. 2(2), 143–160 (2017). doi:10.1016/j.idm.2017.03.002

[16] Guimarães, K.S., Batista, Z.S., Dias, E.L., Guerra, R.M.S.N.C., Costa, A.D.C., Oliveira, A.S., Calabrese, K.S., Cardoso, F.O., Souza, C.S.F., do Vale, T.Z., Gonçalves da Costa, S.C., Abreu-Silva, A.L.: Canine visceral leishmaniasis in São José de Ribamar, Maranhão State, Brazil. Vet. Parasitol. 131(3-4), 305–309 (2005). doi:10.1016/j.vetpar.2005.05.008

[17] Quinnell, R.J., Courtenay, O., Davidson, S., Garcez, L., Lambson, B., Ramos, P., Shaw, J.J., Shaw, M.A., Dye, C.: Detection of Leishmania infantum by PCR, serology and cellular immune response in a cohort study of Brazilian dogs. Parasitology 122(Pt 3), 253–261 (2001)

[18] Felipe, I.M.A., de Aquino, D.M.C., Kuppinger, O., Santos, M.D.C., Rangel, M.E.S., Barbosa, D.S., Barral, A., Werneck, G.L., Caldas, A.d.J.M.: Leishmania infection in humans, dogs and sandflies in a visceral leishmaniasis endemic area in Maranhão, Brazil. Mem. Inst. Oswaldo Cruz 106(2), 207–211 (2011). doi:10.1590/S0074-02762011000200015

[19] Fraga, D.B.M., Solcà, M.S., Silva, V.M.G., Borja, L.S., Nascimento, E.G., Oliveira, G.G.S., Pontes-de-Carvalho, L.C., Veras, P.S.T., Dos-Santos, W.L.C.: Temporal distribution of positive results of tests for detecting Leishmania infection in stray dogs of an endemic area of visceral leishmaniasis in the Brazilian tropics: A 13 years survey and association with human disease. Vet. Parasitol. 190(3-4), 591–594 (2012). doi:10.1016/j.vetpar.2012.06.025

[20] Quinnell, R.J., Carson, C., Reithinger, R., Garcez, L.M., Courtenay, O.: Evaluation of rK39 Rapid Diagnostic Tests for Canine Visceral Leishmaniasis: Longitudinal Study and Meta-Analysis. PLoS Negl. Trop. Dis. 7(1), 1992 (2013). doi:10.1371/journal.pntd.0001992

[21] Werneck, G.L.: Visceral leishmaniasis in Brazil: rationale and concerns related to reservoir control. Rev. Saude Publica 48(5), 851–856 (2014). doi:10.1590/S0034-8910.2014048005615

[22] World Health Organisation: Control of the leishmaniases: Report of a meeting of the who expert committee on the control of leishmaniases. WHO Technical Report Series 949 (2010)

[23] Minnistério da Saúde: Manual de Vigilância e Controle da Leishmaniose Visceral. 1st edition. Brasília: Editora do Ministério da Saúde (2014)

[24] Quinnell, R.J., Dye, C.: Correlates of the peridomestic abundance of Lutzomyia longipalpis (Diptera: Psychodidae) in Amazonian Brazil. Med. Vet. Entomol. 8(3), 219–224 (1994)

[25] Kelly, D.W. and Mustafa, Z. and Dye, C.: Density-dependent feeding success in a field population of the sandfly, Lutzomyia longipalpis. J. Med. Entomol. 65(4), 517–527 (1996)

[26] Alexander, B., Lopes de Carvalho, R., McCallum, H., Pereira, M.H.: Role of the Domestic Chicken (Gallus gallus)in the Epidemiology of Urban Visceral Leishmaniasis in Brazil. Emerg. Infect. Dis. 8(12), 1480–1485 (2002). doi:10.3201/eid0812.010485

[27] Karavadra, S.: Evaluation of Social and Economic Factors Affecting Implementation of Novel Vector Control in Brazil, (2010). MSc thesis, University of Warwick

[28] Courtenay, O., Quinnell, R.J., Garcez, L.M., Shaw, J.J., Dye, C.: Infectiousness in a cohort of brazilian dogs: why culling fails to control visceral leishmaniasis in areas of high transmission. J. Infect. Dis. 186(9), 1314–1320 (2002). doi:10.1086/344312

[29] Quinnell, R.J., Courtenay, O.: Transmission, reservoir hosts and control of zoonotic visceral leishmaniasis. Parasitology 136(14), 1915–1934 (2009). doi:10.1017/S0031182009991156

[30] Quinnell, R.J., Courtenay, O., Garcez, L., Dye, C.: The epidemiology of canine leishmaniasis: transmission rates estimated from a cohort study in Amazonian Brazil. Parasitology 115(Pt 2), 143–156 (1997)

[31] Quinnell, R.J., Dye, C.: An experimental study of the peridomestic distribution of Lutzomyia longipalpis (Diptera: Psychodidae). Bull. Entomol. Res. 84(3), 379–382 (1994). doi:10.1017/S0007485300032508

[32] Courtenay, O., Gillingwater, K., Gomes, P.A.F., Garcez, L.M., Davies, C.R.: Deltamethrin-impregnated bednets reduce human landing rates of sandfly vector Lutzomyia longipalpis in Amazon households. Med. Vet. Entomol. 21(2), 168–176 (2007). doi:10.1111/j.1365-2915.2007.00678.x

[33] Dilger, E.: The effects of host-vector relationships and density dependence on the epidemiology of visceral leishmaniasis. PhD thesis, University of Warwick (2013)

[34] Dye, C., Davies, C.R., Lainson, R.: Communication among phlebotomine sandflies: a field study of domesticated Lutzomyia longipalpis populations in Amazonian Brazil. Anim. Behav. 42(2), 183–192 (1991). doi:10.1016/S0003-3472(05)80549-4

[35] Varma, A., Morbidelli, M., Wu, H.: Parametric Sensitivity in Chemical Systems. Cambridge University Press, Cambridge (1999). doi:10.1017/CBO9780511721779

[36] Damiani, C., Filisetti, A., Graudenzi, A., Lecca, P.: Parameter sensitivity analysis of stochastic models: Application to catalytic reaction networks. Comput. Biol. Chem. 42, 5–17 (2013). doi:10.1016/j.compbiolchem.2012.10.007

[37] Duthie, M.S., Lison, A., Courtenay, O.: Advances toward Diagnostic Tools for Managing Zoonotic Visceral Leishmaniasis. Trends Parasitol. (2018). doi:10.1016/j.pt.2018.07.012

[38] Rock, K.S., Quinnell, R.J., Medley, G.F., Courtenay, O.: Progress in the Mathematical Modelling of Visceral Leishmaniasis. Adv. Parasitol. 94, 49–131 (2016). doi:10.1016/bs.apar.2016.08.001

[39] Rogers, M.E., Bates, P.A.: Leishmania Manipulation of Sand Fly Feeding Behavior Results in Enhanced Transmission. PLoS Pathog. 3(6), 91 (2007). doi:10.1371/journal.ppat.0030091

[40] Stamper, L.W., Patrick, R.L., Fay, M.P., Lawyer, P.G., Elnaiem, D.-E.A., Secundino, N., Debrabant, A., Sacks, D.L., Peters, N.C.: Infection Parameters in the Sand Fly Vector That Predict Transmission of Leishmania major. PLoS Negl. Trop. Dis. 5(8), 1288 (2011). doi:10.1371/journal.pntd.0001288

[41] Cameron, M.M., Acosta-Serrano, A., Bern, C., Boelaert, M., den Boer, M., Burza, S., Chapman, L.A.C., Chaskopoulou, A., Coleman, M., Courtenay, O., Croft, S., Das, P., Dilger, E., Foster, G., Garlapati, R., Haines, L., Harris, A., Hemingway, J., Hollingsworth, T.D., Jervis, S., Medley, G., Miles, M., Paine, M., Picado, A., Poché, R., Ready, P., Rogers, M., Rowland, M., Sundar, S., de Vlas, S.J., Weetman, D.: Understanding the transmission dynamics of Leishmania donovani to provide robust evidence for interventions to eliminate visceral leishmaniasis in Bihar, India. Parasit. Vectors 9, 25 (2016). doi:10.1186/s13071-016-1309-8

[42] Courtenay, O., Peters, N.C., Rogers, M.E., Bern, C.: Combining epidemiology with basic biology of sand flies, parasites, and hosts to inform leishmaniasis transmission dynamics and control. PLOS Pathog. 13(10), 1006571 (2017). doi:10.1371/journal.ppat.1006571

[43] Moreira, E.D., Mendes de Souza, V.M., Sreenivasan, M., Nascimento, E.G., Pontes de Carvalho, L.: Assessment of an optimized dog-culling program in the dynamics of canine Leishmania transmission. Vet. Parasitol. 122(4), 245–252 (2004). doi:10.1016/j.vetpar.2004.05.019

[44] Nunes, C.M., de Lima, V.M.F., de Paula, H.B., Perri, S.H.V., de Andrade, A.M., Dias, F.E.F., Burattini, M.N.: Dog culling and replacement in an area endemic for visceral leishmaniasis in Brazil. Vet. Parasitol. 153(1-2), 19–23 (2008). doi:10.1016/j.vetpar.2008.01.005

[45] Dias, R.A., Baquero, O.S., Guilloux, A.G.A., Moretti, C.F., de Lucca, T., Rodrigues, R.C.A., Castagna, C.L., Presotto, D., Kronitzky, Y.C., Grisi-Filho, J.H.H., Ferreira, F., Amaku, M.: Dog and cat management through sterilization: Implications for population dynamics and veterinary public policies. Prev. Vet. Med. 122(1-2), 154–163 (2015). doi:10.1016/j.prevetmed.2015.10.004

[46] Harhay, M.O., Olliaro, P.L., Costa, D.L., Costa, C.H.N.: Urban parasitology: visceral leishmaniasis in Brazil. Trends Parasitol. 27(9), 403–409 (2011). doi:10.1016/j.pt.2011.04.001

[47] Alexander, B., Lopes de Carvalho, R., McCallum, H., Pereira, M.H.: Role of the Domestic Chicken (Gallus gallus)in the Epidemiology of Urban Visceral Leishmaniasis in Brazil. Emerg. Infect. Dis. 8(12), 1480–1485 (2002). doi:10.3201/eid0812.010485

[48] Bern, C., Courtenay, O., Alvar, J.: Of Cattle, Sand Flies and Men: A Systematic Review of Risk Factor Analyses for South Asian Visceral Leishmaniasis and Implications for Elimination. PLoS Negl. Trop. Dis. 4(2), 599 (2010). doi:10.1371/journal.pntd.0000599

[49] Belo, V.S., Struchiner, C.J., Werneck, G.L., Barbosa, D.S., de Oliveira, R.B., Neto, R.G.T., da Silva, E.S.: A systematic review and meta-analysis of the factors associated with Leishmania infantum infection in dogs in Brazil. Vet. Parasitol. 195(1-2), 1–13 (2013). doi:10.1016/j.vetpar.2013.03.010

[50] Bray, D.P., Bandi, K.K., Brazil, R.P., Oliveira, A.G., Hamilton, J.G.C.: Synthetic Sex Pheromone Attracts the Leishmaniasis Vector Lutzomyia longipalpis (Diptera: Psychodidae) to Traps in the Field. J. Med. Entomol. 46(3), 428–434 (2009). doi:10.1603/033.046.0303

[51] Bray, D.P., Carter, V., Alves, G.B., Brazil, R.P., Bandi, K.K., Hamilton, J.G.C.: Synthetic Sex Pheromone in a Long-Lasting Lure Attracts the Visceral Leishmaniasis Vector, Lutzomyia longipalpis, for up to 12 Weeks in Brazil. PLoS Negl. Trop. Dis. 8(3), 2723 (2014). doi:10.1371/journal.pntd.0002723

[52] Bray, D.P., Alves, G.B., Dorval, M.E., Brazil, R.P., Hamilton, J.G.: Synthetic sex pheromone attracts the leishmaniasis vector Lutzomyia longipalpis to experimental chicken sheds treated with insecticide. Parasit. Vectors 3(1), 16 (2010). doi:10.1186/1756-3305-3-16

[53] David, J.R., Stamm, L.M., Bezerra, H.S., Souza, R.N., Killick-Kendrick, R., Lima, J.W.O.: Deltamethrin-impregnated dog collars have a potent anti-feeding and insecticidal effect on Lutzomyia longipalpis and Lutzomyia migonei. Mem. Inst. Oswaldo Cruz 96(6), 839–847 (2001). doi:10.1590/S0074-02762001000600018

[54] Silva, J.G.D.e., Werneck, G.L., Cruz, M.d.S.P.e., Costa, C.H.N., de Mendonça, I.L.: Infecção natural de Lutzomyia longipalpis por Leishmania sp. em Teresina, Piaúi, Brasil. Cad. Saude Publica 23(7), 1715–1720 (2007). doi:10.1590/S0102-311X2007000700024

[55] Savani, E.S.M.M., Nunes, V.L.B., Galati, E.A.B., Castilho, T.M., Zampieri, R.A., Floeter-Winter, L.M.: The finding of Lutzomyia almerioi and Lutzomyia longipalpis naturally infected by Leishmania spp. in a cutaneous and canine visceral leishmaniases focus in Serra da Bodoquena, Brazil. Vet. Parasitol. 160(1-2), 18–24 (2009). doi:10.1016/j.vetpar.2008.10.090

[56] Reithinger, R., Coleman, P.G., Alexander, B., Vieira, E.P., Assis, G., Davies, C.R.: Are insecticide-impregnated dog collars a feasible alternative to dog culling as a strategy for controlling canine visceral leishmaniasis in Brazil? Int. J. Parasitol. 34(1), 55–62 (2004). doi:10.1016/j.ijpara.2003.09.006

