## Supplementary material for "Spatio-temporal modelling of *Leishmania infantum* infection among domestic dogs: a simulation study and sensitivity analysis applied to rural Brazil"

### Additional File 1

#### Household-level host distributions

The number of each type of host at each household was assigned in each model run by sampling from distributions of host numbers per household. We obtained these distributions by fitting to survey data from the Marajó region collected in July and August of 2010 at 140 households across seven villages; via a questionnaire, data were collected on the number of adults, adolescents, and children resident in the home, as well as the number of dogs and chickens kept at the home [1].

We fit a Poisson distribution to the data for each host; we also fit a negative binomial distribution when the sample variance was not less than the sample mean. The Poisson distribution was initially chosen and fitted because these data are count data; the negative binomial distribution is also a discrete distribution and is considered an alternative to the Poisson distribution for data where the Poisson assumption of the mean being equal to the variance is not appropriate. Distributions were fitted using maximum likelihood estimation via the `poissfit` and `fitdist` functions from the MATLAB® version R2016b Statistics and Machine Learning Toolbox. We compared the fitted Poisson and negative binomial distributions using the Akaike information criterion (AIC) [2]. The AIC estimates the quality of each fitted distribution relative to the other. Specifically, the difference in AIC values between the Poisson and negative binomial models ($\Delta$AIC) indicates the level of support for the Poisson model fit relative to the negative binomial model fit. As a general rule, a larger difference indicates that the Poisson model is less plausible compared to the negative binomial model. Notably, a $\Delta$AIC$\leq 2$ corresponds to the Poisson model having substantial support relative to the negative binomial model, while a $\Delta$AIC$> 10$ indicates no support for the Poisson model [3].

The results of fitting the Poisson and negative binomial distributions (Supplementary Figure 1) determined that where both distributions were fit the negative binomial distribution was preferred as all $\Delta$AIC were greater than 2 (Supplementary Table 1). Visual inspection provided additional support for this outcome (Supplementary Figure 1).

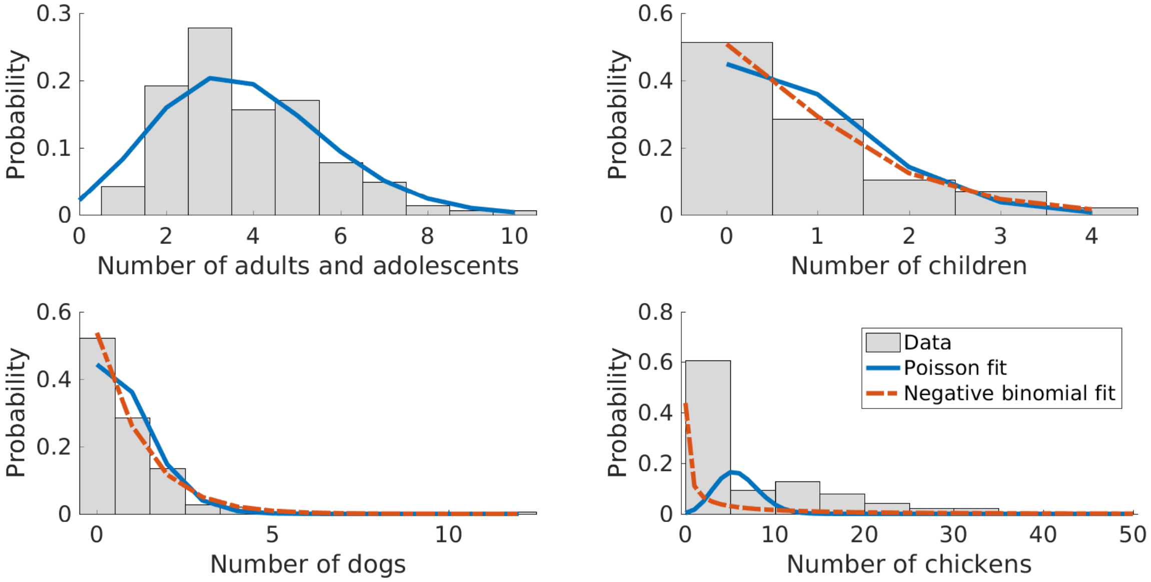


**Supplementary Figure 1: Distributions of the number of hosts per household.** Data (bars), best fit Poisson distributions (blue solid line) and negative binomial distributions (red dashed line, considered when the sample variance was greater than the sample mean) fitted using maximum likelihood estimation for the number of adults and adolescents, children, dogs, and chickens resident at households in Marajó.

**Supplementary Table 1: Poisson and negative binomial model fits for host distributions at households.** For the negative binomial fits, $r$ corresponds to the number of successes and $p$ corresponds to the probability of success. We show that, where fit, the negative binomial distribution was preferred to the Poisson distribution. The parameter maximum likelihood estimates (MLE) for each distribution are reported along with the Akaike information criterion (AIC) for model comparison. MLE estimates are reported to 3 d.p. AIC and $\Delta$AIC values are given to 1 d.p.

| | Poisson fit | | Negative binomial fit | | | |
|---|---|---|---|---|---|---|
| Host type | $\lambda$ MLE | AIC | $r$ MLE | $p$ MLE | AIC | $\Delta$AIC |
| Adults and adolescents | 3.821 | 546.7 | - | - | - | - |
| Children | 0.800 | 351.7 | 2.081 | 0.722 | 347.2 | 4.4 |
| Dogs | 0.814 | 376.6 | 1.206 | 0.597 | 353.1 | 23.4 |
| Chickens | 5.850 | 1816.6 | 0.262 | 0.043 | 712.5 | 1104.1 |

#### Sandfly abundance

Sand fly trapping data from villages in Marajó were used to obtain realistic estimates of the abundance of sand flies, $L_h$, at households. For each household $h$, $L_h$ comprised of two parts: a constant initial estimate $K_h\left(\frac{1}{1-\zeta}\right)$ and a seasonal scaling $v(t)$.

Data on the abundance of female sand flies, specifically the vector species *Lutzomyia longipalpis*, were available from a previous study of 180 households in fifteen villages on Marajó island where sand fly numbers were surveyed using CDC light-traps [4]. The trap-count abundance, $K_h$, was sampled from these data. With traps only capturing a proportion of female sand flies expected at households, a proportion of the female sand fly population, $\zeta$, remained unobserved. Accounting for this inconsistency necessitated the scaling of the trap-count abundance by a factor of $\frac{1}{1-\zeta}$.

Sand fly populations have been observed to exhibit temporal dependencies. To incorporate seasonality into the model, at the beginning of each time step we applied a time-dependent scaling factor, $v(t)$, to all initial abundance estimates. To produce the scaling factor $v(t)$, a smooth trend line was fitted, via a Lowess smoother, to the mean number of female *Lutzomyia longipalpis* trapped over an eight month period across eight different households in the village of Boa Vista, Marajó [5]. The curve was extrapolated over the remaining four months of the year for which no data were available. This highlighted a peak in sand fly abundance during January, at the transition from the dry to wet season (Supplementary Figure 2). Expected vector abundance then dropped and attained its minimum level in May and June, coinciding with the end of the wet season. Normalising the curve between 0 and 1 gave our seasonal scaling factor $v(t)$. Similar temporal patterns were observed in the data split by the eight households (Supplementary Figure 3) and split by location within household (Supplementary Figure 4).

Amalgamating the initial abundance estimate and seasonal scaling components results in the following seasonally-scaled sand fly abundance at household $h$ at time $t$,

$$L_h(t) = K_h\left(\frac{1}{1-\zeta}\right)v(t).$$

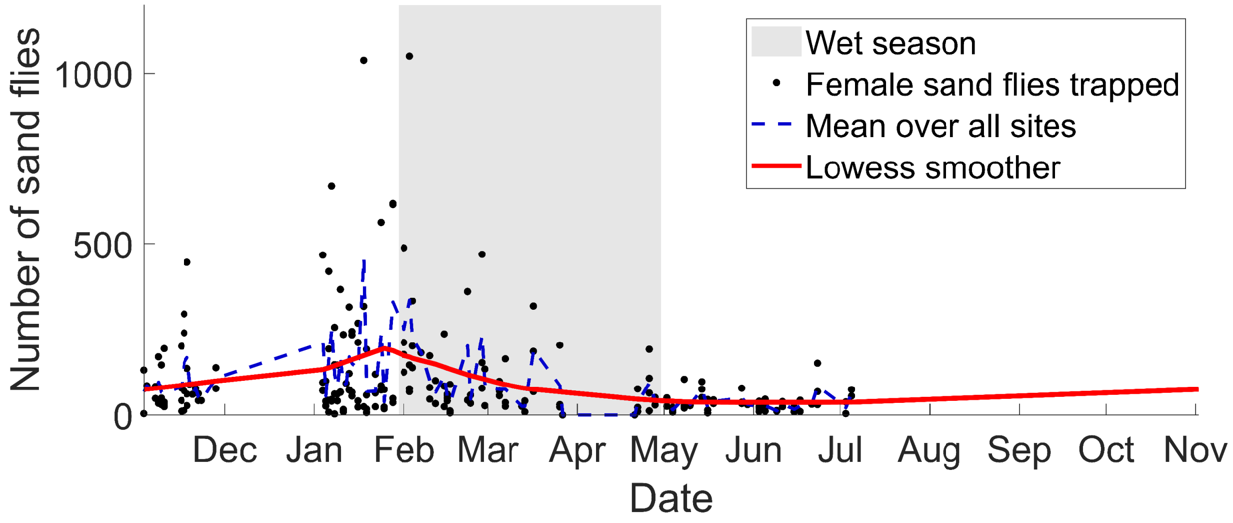


**Supplementary Figure 2: The seasonality of sand fly abundance using Lutzomyia longipalpis data from Marajó.** Data on the number of female *Lutzomyia longipalpis* trapped in a night across eight household sites (blue dots), the mean over household sites in a night (blue line), and a smooth trend line fitted using a Lowess smoother and linearly extrapolated to give values for the remaining four months of the year (red line).

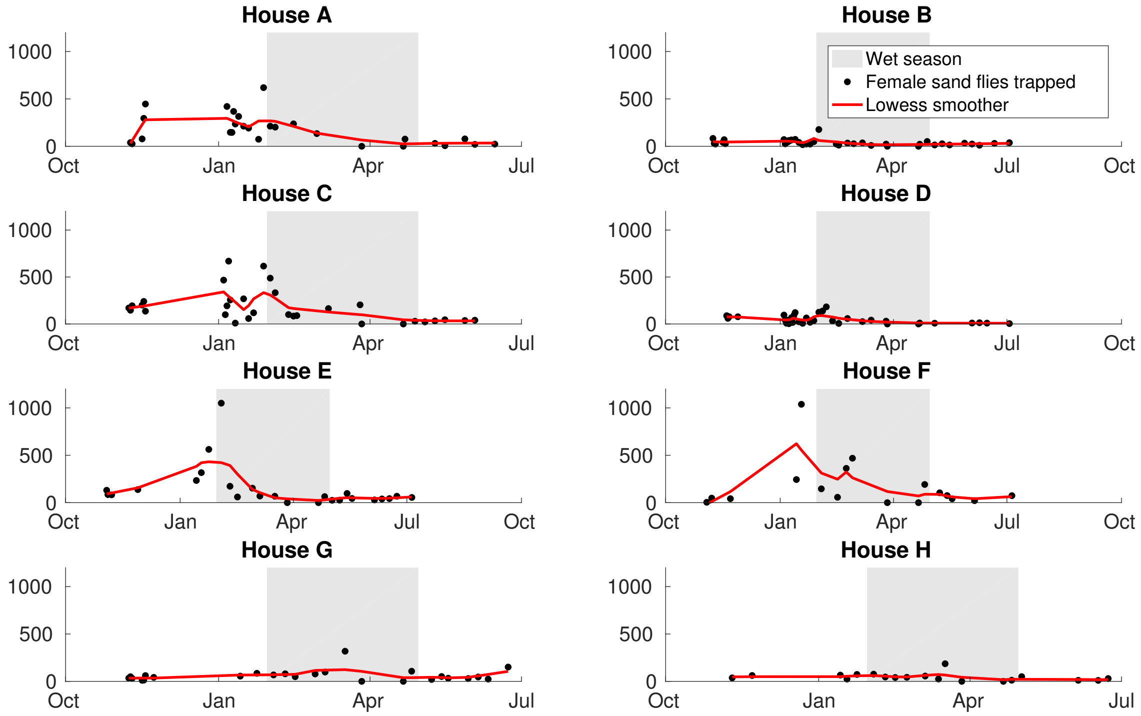


**Supplementary Figure 3: The seasonality of sand fly abundance using data from the Marajó region.** Data on the number of female sand flies trapped in a night in each of the eight household sites (black dots) with a smooth trend line fitted using a Lowess smoother (red line). These households each show similar seasonal patterns.

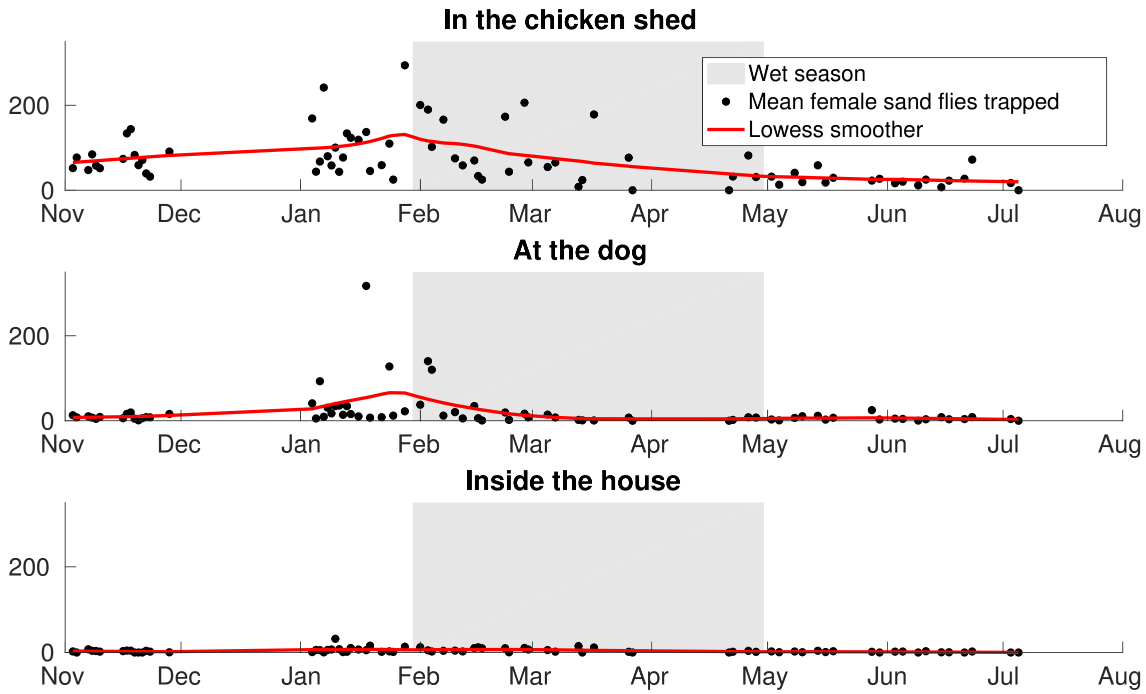


**Supplementary Figure 4: The seasonality of sand fly abundance using data from the Marajó region.** Data on the number of female sand flies trapped in a night at three locations within the trapping sites, presented as the mean over all sites (black dots) with a smooth trend line fitted using a Lowess smoother (red line). The three locations each show similar seasonal patterns.

### Proportion of infectious sand flies

The proportion of infectious vectors at household $h$ was comprised of a time-independent background level of prevalence, $\phi$, constant across all households plus an additional proportion dependent on the number of infectious dogs in the neighbourhood of household $h$. The contribution from each type of infectious dog (high and low infectiousness) was computed separately. We matched the radius $r$ defining this neighbourhood with the maximum sand fly travel distance (taken as 300m at the baseline [6], see Table 1 in main text).

Initially, we computed the proportion of biomass of infectious dogs of type $x$ within radius $r$ of household $h$. This proportion of biomass was subsequently modified by a linear weighting function to account for a reduction in impact with increasing distance from household $h$:

$$B_{h,x}(r,t) = \frac{\sum\limits_{k \in \mathbb{H}_h(r)} \frac{r-d(h,k)}{r} N_{h,x} b_x}{\sum\limits_{k \in \mathbb{H}_h(r)} \frac{r-d(h,k)}{r} \left( \sum\limits_{s \in \text{host type}} N_{h,s} b_s \right)},$$

where $\mathbb{H}_h(r)$ is defined as the set of households within distance $r$ of household $h$, $d(h,k)$ is the distance between households $h$ and $k$, and $N_{h,s}$ is the number of dogs of infectiousness type $s$ at household $h$.

Using these weighted biomass computations for infectious dogs, the proportion of sand flies that were infectious at household $h$ on day $t$ was computed as:

$$\phi_h(t) = \phi + (m_{\text{high}} - \phi)B_{h,d_{\text{high}}}(r,t) + (m_{\text{low}} - \phi)B_{h,d_{\text{low}}}(r,t),$$

where $\phi$ is the constant background level of prevalence, and $m_{\text{high}}$ and $m_{\text{low}}$ are upper bounds on the proportion of infectious sand flies obtained when the only hosts present were high infectiousness dogs or low infectiousness dogs respectively. Under an assumption that 80% of transmission from dogs to sand flies is caused by high infectiousness dogs, with the remaining 20% of total transmission events contributed by infected dogs with low infectiousness [7], the explicit calculations for $m_{\text{low}}$ and $m_{\text{high}}$ were as follows:

$$m_{\text{low}} = \frac{0.2 m_{\text{avg}}}{\tilde{\pi}_{\text{low}}}, \quad m_{\text{high}} = \frac{0.8 m_{\text{avg}}}{\tilde{\pi}_{\text{high}}},$$

with $\tilde{\pi}_{\text{low}}$ and $\tilde{\pi}_{\text{high}}$ denoting the proportion of infectious dogs that have low and high infectiousness, respectively, and $m_{\text{avg}}$ corresponding to the proportion of infectious sand flies obtained when the only hosts present are infectious dogs, obtained by averaging over both high and low infectiousness dogs.

### Sensitivity coefficients

In a stochastic modelling framework, like the one developed in this study, outputs do not take a unique value. Instead, they take a range of values with a given probability, defined by a probability density function $f$. Therefore, to calculate a stochastic sensitivity coefficient for each parameter we followed the procedure outlined in Damiani et al. [8]. In brief, this technique evaluates the sensitivity coefficient $\Upsilon_p^u$ of the output variable of interest $u$ with respect to each parameter $p$,

$$\Upsilon_p^u = \int_{\Omega p} \left\{ \int_{\Omega u} \left| \frac{\partial f(u(p))}{\partial p} \right| f(u(p))\, du \right\} dp, \tag{1}$$

where $\Omega u$ is the domain of integration of $u$. Due to the computational demands of evaluating the density function for the entire parameter space, the integrals in Equation 1 were calculated on a finite domain. The probability density function $f(u(p))$ and the partial derivatives $\frac{\partial f(u(p))}{\partial p}$ were estimated using non-parametric kernel methods using simulation outputs from the model.

### References


[1] Karavadra, S.: Evaluation of social and economic factors affecting implementation of novel vector control in Brazil. MSc thesis, University of Warwick, 2010.

[2] Akaike, H.: A new look at the statistical model identification. IEEE Transactions on Automatic Control, 19:6(716–723), 1974.

[3] Burnham, K.P., Anderson, D.R.: Model Selection and Multimodel Inference: A Practical Information-Theoretic Approach. Springer New York, NY, 2004.

[4] Quinnell, R.J. and Dye, C.: Correlates of the peridomestic abundance of Lutzomyia longipalpis (Diptera: Psychodidae) in Amazonian Brazil.. Medical and Veterinary Entomology, 8:3(219–224), 1994.

[5] Dilger, E.: The effects of host-vector relationships and density dependence on the epidemiology of visceral leishmaniasis. PhD thesis, University of Warwick, 2013.

[6] Dye, C., Davies, C.R. and Lainson, R.: Communication among phlebotomine sandflies: a field study of domesticated Lutzomyia longipalpis populations in Amazonian Brazil. Animal Behaviour, 42:2(183–192), 1991.

[7] Courtenay, O., Quinnell, R.J., Garcez, L.M., Shaw, J.J. and Dye, C.: Infectiousness in a cohort of brazilian dogs: why culling fails to control visceral leishmaniasis in areas of high transmission. The Journal of Infectious Diseases, 186:9(1314–1320), 2002.

[8] Damiani, C., Filisetti, A., Graudenzi, A., Lecca, P.: Parameter sensitivity analysis of stochastic models: Application to catalytic reaction networks. Computational Biology and Chemistry, 42(5–17), 2013.
