## Supplementary material for "Spatio-temporal modelling of *Leishmania infantum* infection among domestic dogs: a simulation study and sensitivity analysis applied to rural Brazil"

#### Abstract

**Background:** The parasite *Leishmania infantum* causes zoonotic visceral leishmaniasis (VL), a potentially fatal vector-borne disease of canids and humans. Zoonotic VL poses a significant risk to public health, with regions of Latin America being particularly afflicted by the disease.

*Leishmania infantum* parasites are transmitted between hosts during blood feeding by infected female phlebotomine sand flies. With ~~domestic dogs being~~ a principal reservoir host of ~~*Leishmania*~~ L. *infantum*~~, a primary focus of research efforts has been to understand disease transmission dynamics among dogs. The intention being that~~ being domestic dogs, limiting prevalence in this reservoir ~~will~~ may result in a reduced risk of infection for the human population. To this end, a primary focus of research efforts has been to understand disease transmission dynamics among dogs. One way this can be achieved is through the use of mathematical models. ~~.~~

~~A Poisson distribution was fitted to the data for each host; a negative binomial distribution was also fitted when the sample variance was less than the sample mean. Distributions were fitted using maximum likelihood estimation via the poissfitand fitdistfunctions from the MATLAB® Statistics and Machine Learning Toolbox. Fitted Poisson and negative binomial distributions were compared using the Akaike information criterion (AIC) [28].~~

#### Infection progression in dogs

The natural history of *L. infantum* infection in dogs consists of susceptible and infected states. Prior work has established heterogeneities in the infectiousness of dogs (transmission of *L. infantum* to the vector) [2, 29, 30]. This was represented in the model by stratifying Specifically, this heterogeneity in infectiousness results in infected states representing highly infectious dogs (responsible for 80% or greater of all observed transmission events), mildly infectious dogs (contributing to 20% or less of total transmission events), and noninfectious dogs that, although infected, never transmit the *L. infantum* parasite back to susceptible sand flies [29].

For modelling purposes we therefore stratified infected dogs into four states: (i) latently infected; (ii) never infectious; (iii) low infectiousness; (iv) high infectiousness (Figure 3). Particularly noteworthy is that never infectious dogs, although infected, do not transmit the *L. infantum* parasite back to susceptible sand flies. Susceptible dogs became latently infected at a rate dependent on the force of infection$\lambda$; full details of this will follow. Movement between the latently infected state and the remaining three infected states occurred at constant rates. Note that a fully recovered state was not included as the complete cure of *L. infantum* infected dogs is rare (even after treatment), validated by experimental observations finding minimal seroreversion from *L. infantum* parasite seropositivity [31].

Deaths could occur from every state in the model and the mortality rates were state specific ($\mu_{\text{Sus}}$, $\mu_{\text{NeverInf}}$, $\mu_{\text{LowInf}}$, $\mu_{\text{HighInf}}$)differed between states. Upon death from any state, a new dog was introduced into the same household at a given replacement rate($1/\psi$). These newly-introduced . Newly-introduced dogs were placed either in the susceptible state or one of the infected states(with probabilities $1-\xi$ and $\xi$ accordingly), encapsulating both birth and immigration into the study region. It follows that the initially sampled initial dog populations corresponded to the maximum attainable dog population size per household.

#### Force of infection

Sand fly dynamics operate on a faster time-scale compared to the other host species and processes considered in the model; sand flies have an estimated life expectancy of a number of weeks at most [10]. For that reason, we did not explicitly track the transitions of sand flies between the susceptible and infectious states at an individual level. We instead considered sand fly populations at each house as a collective which exert a force of infection, $\lambda$, on dogs at household $h$ at time $t$ in the following way,

$$\lambda_h(t) = \alpha \times \delta \times L_h(t) \times \eta_{h,\text{dog}}(t) \times \phi_h(t), \qquad (1)$$

where $\alpha$ is the biting rate of sand flies, $\delta$ is the probability of *L. infantum* transmission to dogs as a result of a single bite from an infectious sand fly, $L_h$ is the abundance of sand flies at household $h$, $\eta_{h,\text{dog}}$ is the host preference of sand flies towards dogs (that is, the probability of sand flies biting dogs at household $h$ as opposed to any other host), and $\phi_h$ is the proportion of sand flies that are infectious at household $h$.

As most sand fly activity occurs in the evening when most the majority of hosts will be within their household [32, 33], we discretised our simulations into daily time steps. This Using daily time steps gave the following probability for a susceptible dog at household $h$ to become infected on day $t$:

$$p_h(t) = 1 - e^{-\lambda_h(t)}. \qquad (2)$$

The biting rate and probability of an infected sand fly transmitting *L. infantum* to a dog as a result of a single bite were constant in the model. In contrast, sand fly abundance, host preference, and the proportion of sand flies infected at each household were time-dependent; we now outline the computation of each time-dependent component.

**Sandfly abundance:** Sand fly trapping data from villages in Marajó were used to obtain realistic estimates of the abundance of sand flies, $L_h$, at households. ~~For each household $h$,~~ As sand fly populations have been observed to exhibit temporal dependencies $L_h$ comprised of two parts: a constant initial estimate ~~$K_h \left(\frac{1}{1-\zeta}\right)$~~ and a seasonal scaling ~~$v(t)$~~factor.

Data on the abundance of female sand flies, specifically the vector species *Lutzomyia longipalpis*, were available from a previous study of 180 households in fifteen villages on Marajó island where sand fly numbers were surveyed using CDC light-traps [24]. The ~~trap-count abundance , $K_h$,~~ constant initial estimate of abundance was sampled from these data ~~. With traps only capturing a proportion of~~ and scaled by the expected proportion of unobserved female sand flies ~~expected at households, a proportion of the female sand fly population, $\zeta$, remained unobserved. Accounting for this inconsistency necessitated the scaling of the trap-count abundance by a factor of $\frac{1}{1-\zeta}$~~ at households.

~~Sand fly populations have been observed to exhibit temporal dependencies. To incorporate this seasonality into the model, we applied a time-dependent scaling factor, $v(t)$, to all initial abundance estimates at the beginning of each time step. To produce the scaling factor $v(t)$, a smooth trend line was fitted, via a Lowess smoother, to the~~ Data on the mean number of female *Lutzomyia longipalpis* trapped over an eight month period across eight different households in the village of Boa Vista, Marajó [34] ~~. The curve was extrapolated over the remaining four months of the year for which no data were available and then normalised by dividing by the maximum value.~~

~~Putting this together leads to the following seasonally-scaled~~ were then used to find the seasonal scaling factor. Full details of this procedure to estimate sand fly abundance ~~at household $h$ at time $t$,~~

$$L_h(t) = K_h \left(\frac{1}{1-\zeta}\right) v(t).$$

can be found in Additional File 1.

**Host preference:** To parameterise sand fly biting preference towards the host species of interest, we drew on findings from field and laboratory experiments performed in this setting by Quinnell et al [7]. These experiments concluded that the attractiveness of the three host species ~~under study~~ we consider (humans, dogs and chickens) to the *Lutzomyia longipalpis* vector seemed to largely be a function of the relative host sizes.

These experimental findings were used to allocate a portion of sand fly bites to each host type at each household, via each host type being assigned the following biomass value relative to chickens:

- 1 dog = 2 chickens,
- 1 child = 5 chickens,
- 1 adult or adolescent = 10 chickens (using adult-child ratio: 1 adult = 2 children).

The preference, $\eta_{h,x}$, towards host type $x$ at household $h$ was computed as a simple proportion of the total biomassas follows,

$$\eta_{h,x}(t) = \frac{N_{h,x} b_x}{\sum\limits_{s \in \text{host type}} N_{h,s} b_s}, \tag{3}$$

where $N_{h,x}$ is the number of host type $x$ at household $h$ and $b_x$ is the biomass of host type $x$ relative to chickens. So, for example, $b_{\text{dog}} = 2$.

**Proportion of infectious sandfliessand flies:** The proportion of infectious vectors at household $h$ was comprised of a time-independent background level of prevalence , $\phi$, that was constant across all households plus and an additional proportion dependent on the number of infectious dogs in the neighbourhood of household $h$. The contribution from each type of infectious dog (high and low infectiousness) was computed separately. We matched the radius $r$ We informed the radius defining this neighbourhood with by matching it to the maximum sand fly travel distance (taken as 300m at the baseline with a range from 20m to 2km to fully explore the parameter space [35], see Table 1).

Initially, we computed the proportion of biomass of infectious dogs of type $x$ within radius $r$ of household $h$. This proportion of biomass was then weighted using a linear weighting function to account for the reduction in impact with increasing distance from household $h$: where $\mathbb{H}_h(r)$ is defined as the set of households within distance $r$ of household $h$, $d(h,k)$ is the distance between households $h$ and $k$, and $N_{h,s}$ is the number of dogs of infectiousness type $s$ at household $h$.

Using these weighted biomass computations for infectious dogs, the proportion of sand flies that were infectious at household $h$ on day $t$ was computed as:

$$\phi_h(t) = \phi + (m_{\text{high}} - \phi) B_{h,d_{\text{high}}}(r,t) + (m_{\text{low}} - \phi) B_{h,d_{\text{low}}}(r,t),$$

where $\phi$ is the constant background level of prevalence, and $m_{\text{high}}$ and $m_{\text{low}}$ are upper bounds on the proportion of infectious sand flies obtained when the only hosts present were high infectiousness dogs or low infectiousnessdogs respectively. These quantities were obtained The contribution from each type of infectious dog (high and low infectiousness) was computed separately under an assumption that 80% of transmission from dogs to sand flies is caused by high infectiousness dogs [29].

The explicit calculations for $m_{\text{low}}$ and $m_{\text{high}}$ were as follows, with $\tilde{\pi}_{\text{low}}$ and $\tilde{\pi}_{\text{high}}$ denoting the proportion of infectious dogs that have low and high infectiousness, respectively:

$$m_{\text{low}} = \frac{0.2 m_{\text{avg}}}{\tilde{\pi}_{\text{low}}}, \quad m_{\text{high}} = \frac{0.8 m_{\text{avg}}}{\tilde{\pi}_{\text{high}}},$$

where $m_{\text{avg}}$ corresponds to , with the remaining 20% of total transmission events contributed by infected dogs with low infectiousness [29]. Further details on our calculation of the proportion of infectious sand flies obtained when the only hosts present are infectious dogs, obtained by averaging over both high and low infectiousness dogs. sand flies that were infectious are given in Additional File 1.

#### 2.2 Model outputs

Being a stochastic model, the infection dynamics vary on separate simulation runs even with all parameters and other model inputs remaining fixed. By running the model multiple times we obtain an ensemble of model outputs. This collection of model outputs permits the calculation of a variety of summary statistics describing the epidemiology of *L. infantum* infection among domestic dogs, such as prevalence and incidence.

We focus here on the prevalence of infection. To clarify, an infection case refers to any dog harbouring *L. infantum* parasites, including those with and without canine VL symptoms. Thus, we defined infection prevalence at time $t$ as the aggregated percentage of dogs in the latently infected, never infectious, low infectiousness and high infectiousness states, which is equivalent to calculating the proportion of dogs not in the susceptible state:

$$\text{prevalence}(t) = \frac{\text{\# of dogs in population - \# of dogs in susceptible state}}{\text{\# of dogs in population}} \times 100. \tag{4}$$

The daily prevalence estimates were used to obtain an average prevalence, ~~which was~~ defined as the mean of the daily prevalence estimates in a specified time period. Throughout this work, all average prevalence values were computed from the daily prevalence values over the final year (365 days) of each simulation run. Mathematically, with $T$ denoting maximum time, ~~this~~ average infection prevalence may be expressed as

$$\text{Average infection prevalence} = \frac{\sum_{t=T-364}^{T} \text{prevalence}(t)}{365}. \tag{5}$$

#### 2.3 Model summary

In summary, the arrangement of and interaction between the individual pieces of our stochastic, spatial, individual-based model for *L. infantum* infection dynamics in dogs are displayed in Figure 4. We refer to the process in Figure 4 as one run of the simulation.

#### 2.4 Sensitivity analysis

##### Parameter values

We carried out a sensitivity analysis to determine the robustness of the model behaviour to the biological parameter values and to ascertain which parameters had a high impact on the average prevalence as predicted by the model. The values tested for each parameter were within plausible ranges informed via published estimates from the literature and unpublished fieldwork data (Table 1).

We undertook a one-at-a-time sensitivity analysis. That is, each parameter was varied in turn while all others remained at their baseline value. We considered 46 parameter sets (Table 1), and for each individual parameter set we performed 1000 separate model simulation runs. The elapsed simulation time in each run corresponded to ten years.

##### Sensitivity coefficients

In addition to comparing the changes in average prevalence given by each parameter set, we computed sensitivity coefficients. These reflect the ratios between the size of the change in a ~~biological model output and the perturbation of system parameters that cause this change~~ model output (in this case, the change in average VL prevalence) with the corresponding size of the change in the parameter [36]. ~~However, outputs do not take a unique value~~ The sensitivity coefficients therefore account for the different ranges in the values tested for each parameter (Table 1) and ensure that the parameters can be sensibly compared.

However, in a stochastic modelling framework~~. Instead, they take a range of values with a given probability, defined by a probability density function $f$.~~

~~Therefore, to calculate a stochastic sensitivity coefficient for each parameter we followed the procedure~~ , such as this one, model outputs do not take a unique value. To account for stochastic fluctuations while still allowing us to critically analyse the sensitivity of the model parameters, we therefore calculated stochastic sensitivity coefficients (as outlined in Damiani et al. [37]~~. In brief, this technique evaluates the sensitivity coefficient $\Upsilon_p^u$ of the output variable of interest $u$ with respect to each parameter $p$~~,

$$\Upsilon_p^u = \int_{\Omega p} \left\{ \int_{\Omega u} \left| \frac{\partial f(u(p))}{\partial p} \right| f(u(p))\, du \right\} dp,$$

~~where $\Omega u$ is the domain of integration of $u$. Due to the computational demands of evaluating the density function for the entire parameter space, the integrals in **??** were calculated on a finite domain. The probability density function $f(u(p))$ and the partial derivatives $\frac{\partial f(u(p))}{\partial p}$ were estimated using non-parametric kernel methods using simulation outputs from the model.~~

~~We then~~ comprehensive explanation in Additional File 1). We ranked the parameters according to the stochastic sensitivity coefficients, with a larger sensitivity coefficient corresponding to a parameter with higher sensitivity of average VL prevalence to its variation.

All ~~calculations and simulations were carried out~~ simulations were performed in MATLAB® ~~.~~ versions R2014a to R2015a. All other computations and plots were carried out in MATLAB® version R2016b or later.

#### 3 Results

##### 3.1 ~~Curating data~~

###### ~~Household-level host distributions~~

~~We fit distributions to the data on the number of hosts in rural Brazilian households (Figure 2). For the datasets fit with both Poisson and negative binomial distributions, AIC calculations determined the negative binomial distribution to be preferred (Additional file 1: Supplementary Table 1).~~

###### ~~Sandfly seasonality~~

~~Fitting a Lowess smoother to the longitudinal data on female *Lutzomyia longipalpis* capture numbers and extrapolating over the remainder of the year where no data were available highlighted a peak in January at the transition from the dry to wet season (??). Expected vector abundance then dropped and attained its minimum level in May and June, coinciding with the end of the wet season. Normalising this curve between 0 and 1 gave our seasonal scaling factor $v(t)$.~~

~~Similar temporal patterns were observed in the data split by the eight households (Additional file 1: Supplementary Figure 1) and split by location within household (Additional file 1: Supplementary Figure 2).~~

Four parameters associated with sand flies were among the top six ~~parameters in the sensitivity ranking~~most sensitive parameters (Figure 7). The only sand fly-associated parameter ~~absent was~~ that was not among these top six most sensitive parameters was the probability of a susceptible sand fly becoming infected when biting an infectious dog (parameter ID 14)~~(Figure 7)~~.

#### Sensitivity of *L. infantum* infection to biological parameter variation

Running model simulations using baseline biological parameter values set within plausible ranges determined from the literature generated infection prevalence predictions that were within the range of empirical estimates from ~~this region~~ endemic regions of Brazil [16–20]. Variation in infection estimates ~~are~~ is expected as ultimately their precision depends on the ~~sensitivity and specificity of diagnostic tests, the type of test~~ type of diagnostic test used (e.g. molecular vs. immunological), diagnostic test sensitivity and specificity, the choice of clinical sample, and the stage of infection progression [17, 19, 20, 38]. Thus, for example, as dogs acquire parasitological infection prior to detection of serum containing anti-*Leishmania* specific antibodies (seroconversion), seroprevalence data may underestimate true infection rates.

**Visualisation:** EBJ, EH

**Writing - original draft:** EBJ, EH

**Writing - review & editing:** EBJ, EH, SD, ED, OC

All authors read and approved the final version of the manuscript.

#### Financial disclosure

The study was supported by a Wellcome Trust Strategic Translation Award (WT091689MF). The funders had no role in study design, data collection and analysis, decision to publish, or preparation of the manuscript.

### Figures

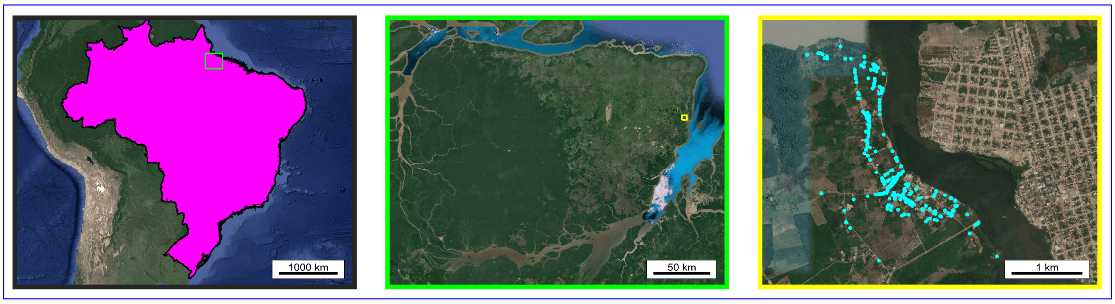


**Figure 1: Locator maps. (Left)** ~~Locator map~~ Map depicting ~~the location of~~ Marajó, situated inside the light green box, within Brazil (shaded in magenta). **(Centre)** ~~Locator map~~ Map depicting ~~the location of~~ Calderao village, situated inside the yellow box, within Marajó. **(Right)** Household locations within Calderao village (cyan filled circles). All map data are from Google and plotted in MATLAB®.

~~**Distributions of the number of hosts per household.** Data (bars), best fit Poisson distributions (blue solid line) and negative binomial distributions where the sample variance was less than the sample mean (red dashed line) fitted using maximum likelihood estimation for the number of adults and adolescents, children, dogs, and chickens resident at households in Marajó.~~

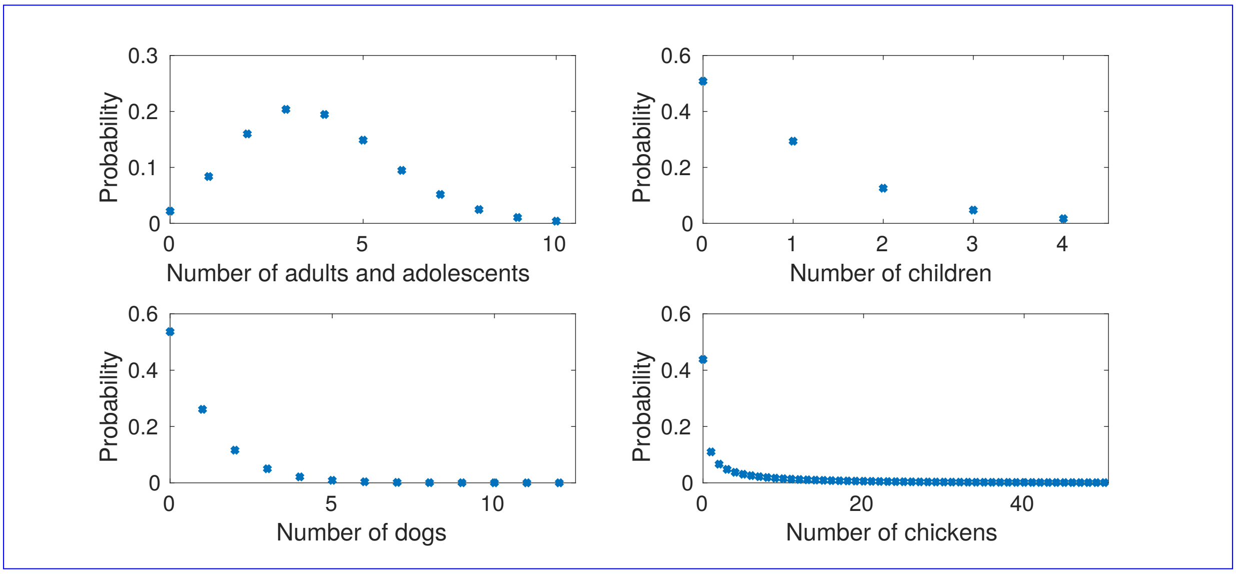


**Figure 2: Distributions of the number of hosts per household. (Top left)** Number of adults and adolescents; **(Top right)** children; **(Bottom left)** Number of dogs; **(Bottom right) Number of chickens.** Full details on how these distributions were obtained can be found in Additional File 1.

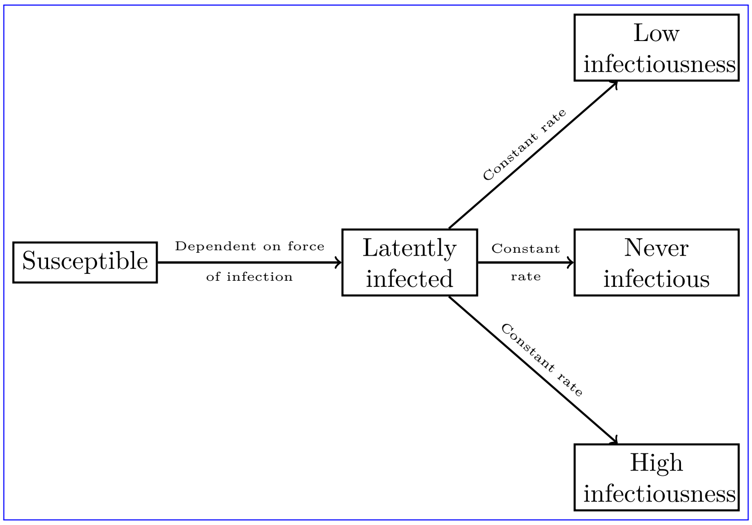


**Figure 3: Model of *L. infantum* infection status in dogs.** Death and replacement of deceased dogs (through birth and immigration) are not shown in the figure.

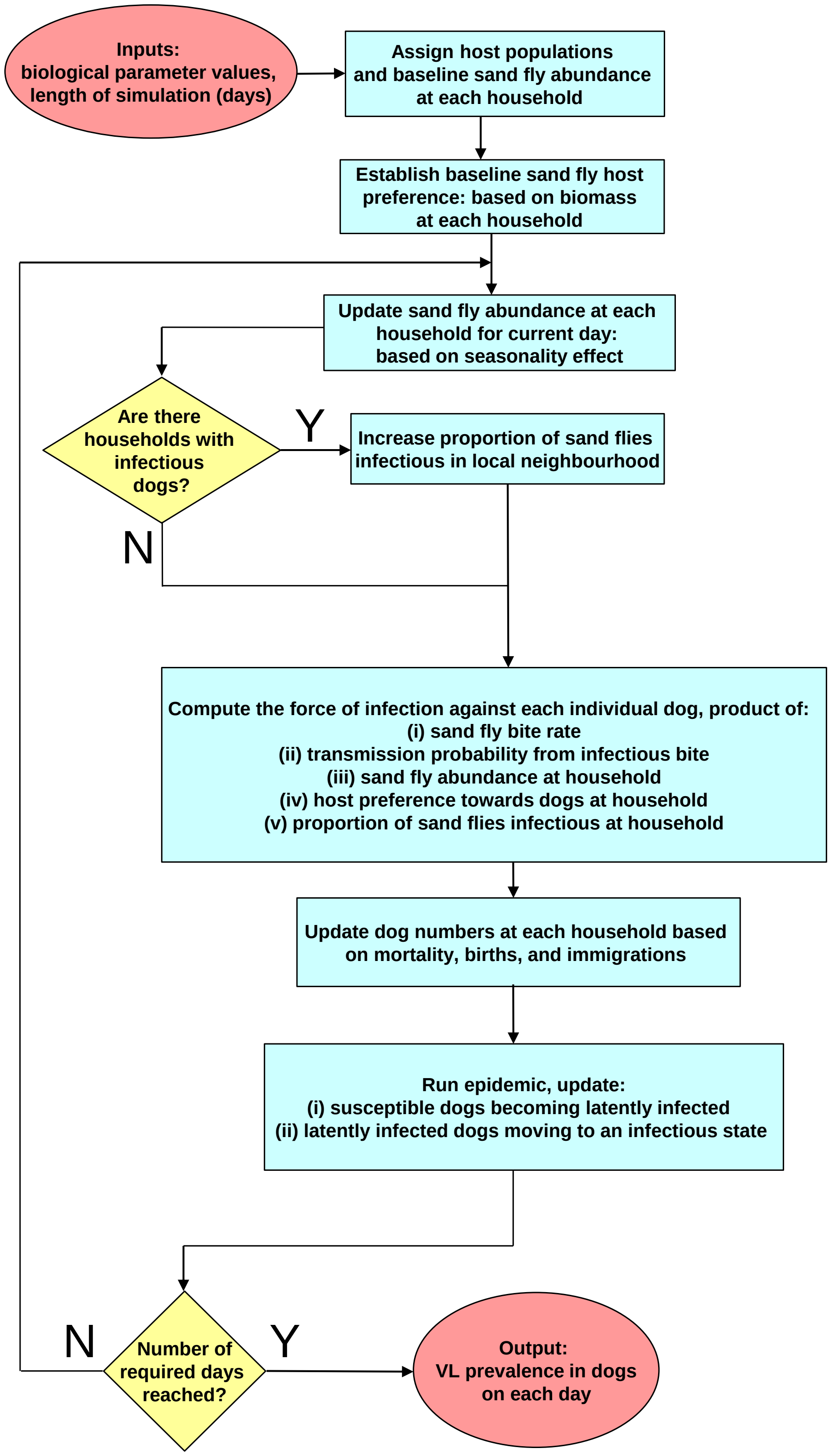


**Figure 4: Visual schematic of model framework for each simulation run.** Red filled ovals represent model inputs and outputs; blue filled rectangles represent actions; yellow filled diamonds represent decisions.

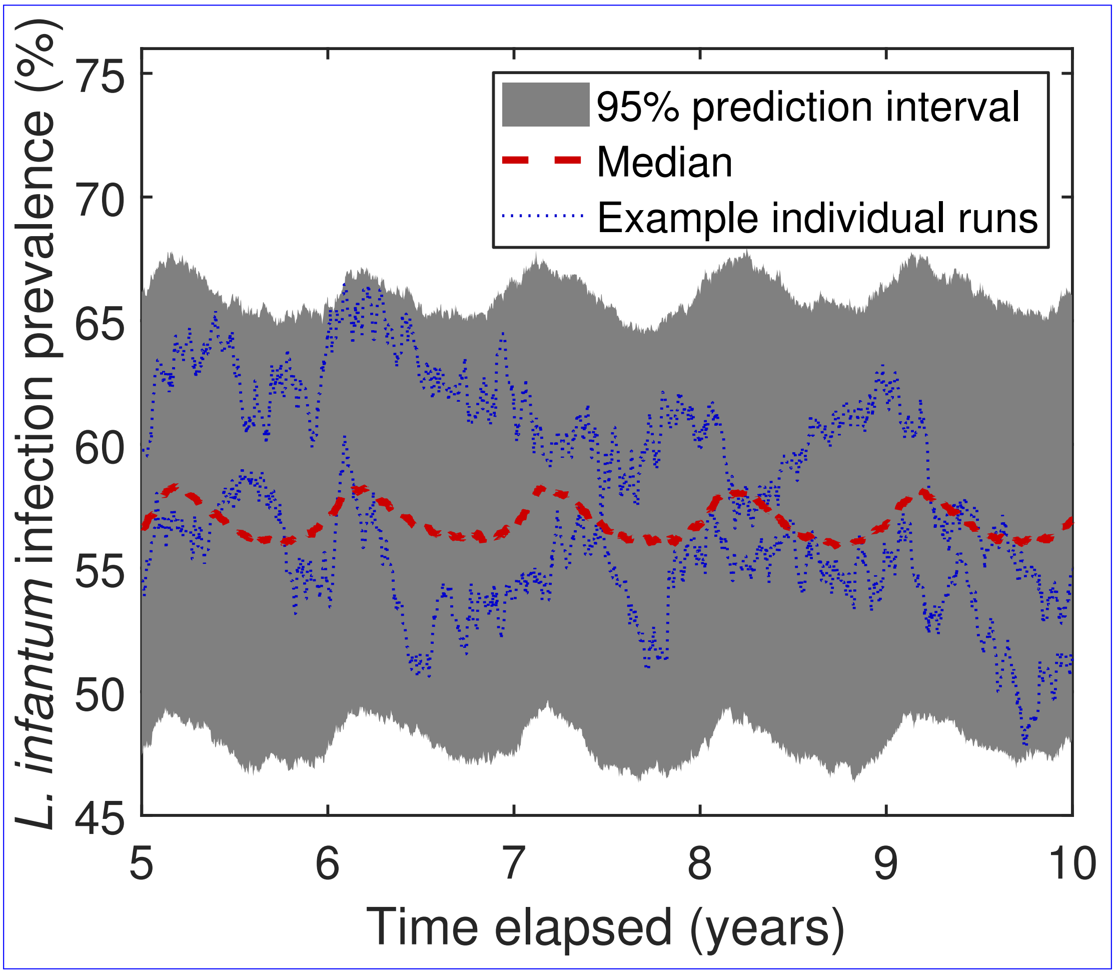


**Figure 5:** ~~**The seasonality of sand fly abundance using data from Marajó.** Data on the number of female sand flies trapped in a night across eight household sites (blue dots)~~**Simulated daily prevalence in domestic dogs using baseline biological parameters.** Dashed, ~~the mean over household sites in a night (blue~~ red line ~~), and a smooth trend line fitted using a Lowess smoother and linearly extrapolated~~ corresponds to ~~give values for~~ the ~~remaining four months of~~ median prevalence and the ~~year (red line)~~grey, filled region depicts the 95% prediction interval at each timestep obtained from 1000 simulation runs. Blue, dotted lines correspond to measured prevalence from two individual simulation runs.

~~**Simulated infection prevalence in domestic dogs using baseline biological parameters.** Dashed, red line corresponds to the median prevalence and the grey, filled region depicts the 95% prediction interval at each timestep obtained from 1000 simulation runs. Blue, dotted lines correspond to measured prevalence from two, individual simulation runs.~~

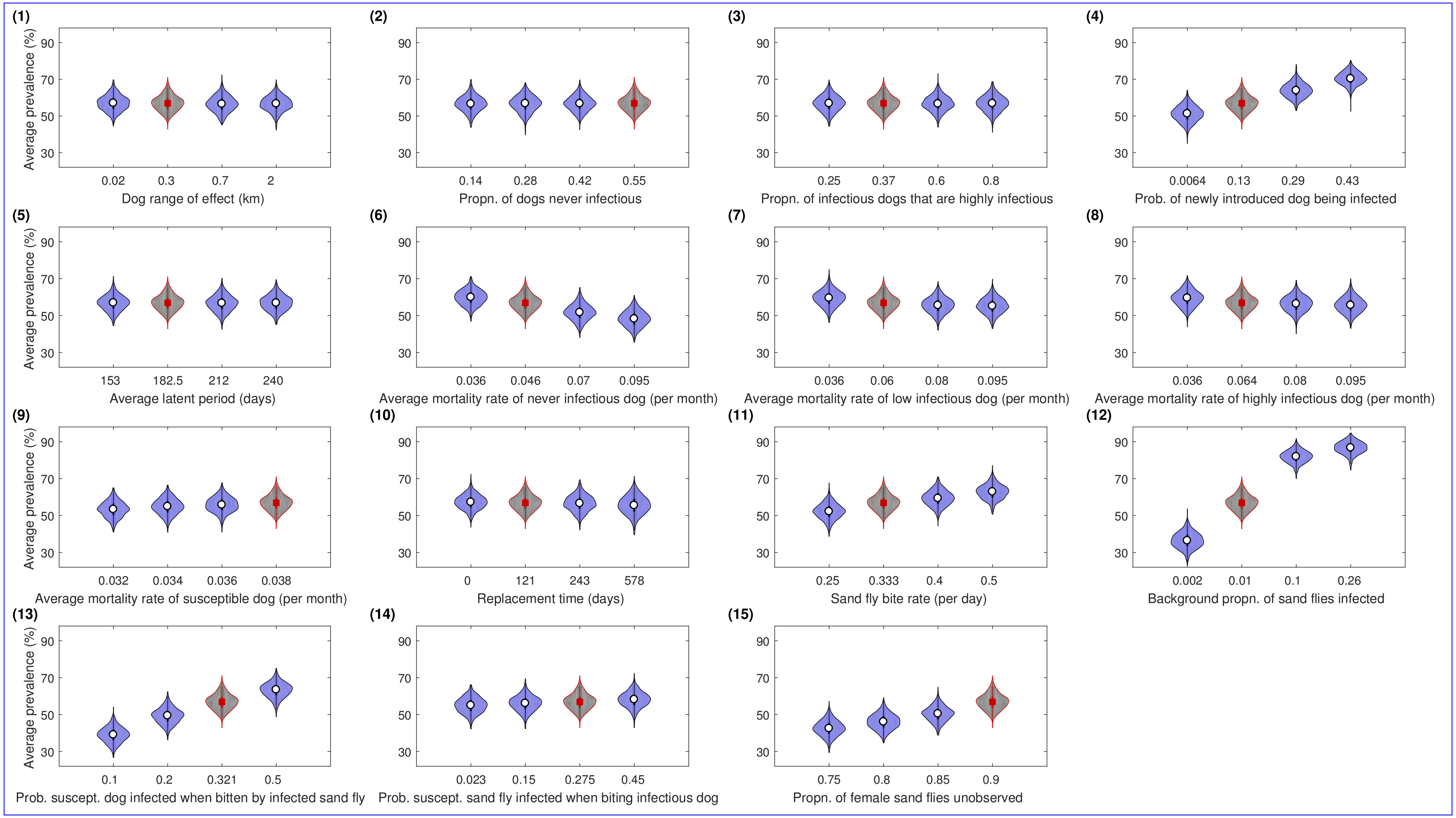


**Figure 6: Violin plots for average infection prevalence under each biological parameter set.** Panel numbering aligns with the parameter ID numbers in Table 1. The average infection prevalence ~~calculation entailed averaging~~ was calculated by the daily prevalence values over the final year of each simulation run. For each parameter set, predicted average infection prevalence distributions were acquired from 1000 simulation runs. The violin plot outlines illustrate kernel probability density, i.e. the width of the shaded area represents the proportion of the data located there. For parameter sets corresponding to the use of the baseline parameter set, violin plot regions are shaded grey with estimated median values represented by a red square. In all other instances, violin plot regions are shaded blue with median values depicted by a white circle.

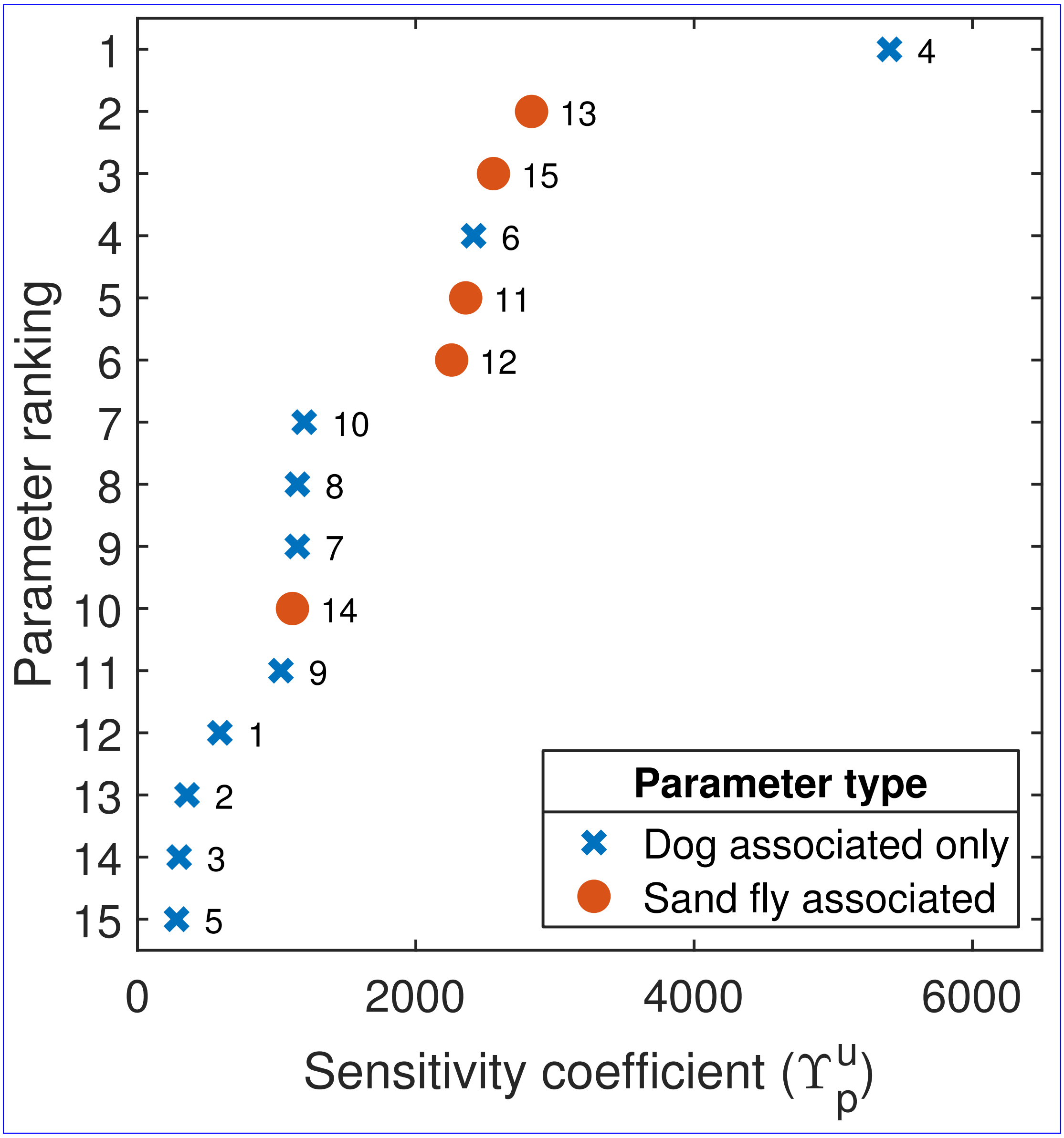


**Figure 7: Stochastic sensitivity coefficient parameter ranking.** The parameter ID linked to each stochastic sensitivity coefficient is placed aside the data point. Blue crosses denote those biological parameters associated with dogs. Filled orange circles correspond to biological parameters associated with ~~sandflies~~sand flies. Average infection prevalence was most sensitive to parameter ID 4 (probability of a newly introduced dog being infected).

### Tables

**Table 1:** Description of measurable biological variables that are used to inform parameters (either directly or after performing additional calculations) in the model. Source listed as OC denotes (O. Courtenay, unpublished observations).

| Param. ID | Symbol | Description | Baseline value | Other values tested | Sources |
|---|---|---|---|---|---|
| 1 | $r$ | Interaction range of dogs (km). | 0.30 | 0.02, 0.7, 2 | [35] |
| 2 | $\pi_{\text{never}}$ | Proportion of infected dogs that are never infectious. | 0.55 | 0.14, 0.28, 0.42 | [29, 30] |
| 3 | $\tilde{\pi}_{\text{high}}$ | Proportion of infectious dogs that are highly infectious. | 0.37 | 0.25, 0.60, 0.80 | [2] |
| 4 | $\xi$ | Probability of a newly introduced dog being infected. | 0.130 | 0.0064, 0.29, 0.43 | [44] |
| 5 | $\nu$ | Per capita rate of progression of dogs from latently infected to a further state (days$^{-1}$). $1/\nu$ is the average duration of the latent period (days). | 0.0055 | 0.0042, 0.0047, 0.0065 | [29] |
| 6 | $\mu_{\text{NeverInf}}$ | Per capita mortality rate for latently infected and never infectious dogs (days$^{-1}$). | 0.0015 | 0.0012, 0.0023, 0.0031 | OC |
| 7 | $\mu_{\text{LowInf}}$ | Per capita mortality rate for dogs with low infectiousness (days$^{-1}$). | 0.0020 | 0.0012, 0.0026, 0.0031 | OC |
| 8 | $\mu_{\text{HighInf}}$ | Per capita mortality rate for dogs with high infectiousness (days$^{-1}$). | 0.0021 | 0.0012, 0.0026, 0.0031 | OC |
| 9 | $\mu_{\text{Sus}}$ | Per capita mortality for susceptible dogs (days$^{-1}$). | 0.00125 | 0.00105, 0.00112, 0.00118 | OC |
| 10 | $\psi$ | Average time (days) for deceased dog to be replaced. | 121 | 0, 243, 578 | [45] |
| 11 | $\alpha$ | Biting rate of sand flies (per day). (Number of times one sand fly would want to bite a host per unit time, if hosts were freely available). | 0.333 | 0.25, 0.40, 0.50 | [35] |
| 12 | $\phi$ | Background proportion of sand flies that are infected. | 0.010 | 0.002, 0.100, 0.260 | [18, 55, 56] |
| 13 | $\delta$ | Probability of *Leishmania* transmission from an infectious sand fly to a susceptible dog given that a contact bite occurs. | 0.321 | 0.10, 0.20, 0.50 | [57] |
| 14 | $m_{\text{avg}}$ | Probability of *Leishmania* transmission from an infectious dog to a susceptible sand fly given that a contact between the two occurs. | 0.275 | 0.023, 0.150, 0.450 | [29] |
| 15 | $\zeta$ | Proportion of female sand fly population not observed in trapping studies. | 0.90 | 0.75, 0.80, 0.85 | [35] |
